# Spiders bring new insight into the eco-evolutionary drivers of body size variation and sexual size dimorphism in arthropods

**DOI:** 10.1101/2024.10.30.621136

**Authors:** Camille Ameline, Denis Lafage, Philippe Vernon, Mickaël Hedde, Julien Pétillon

## Abstract

Body size has been used thoroughly in arthropod ecology as a reliable trait to assess fitness responses to changes in environmental factors. Among these, spiders represent a large and diverse group, colonizing almost all terrestrial habitats. Here, we propose a review on intraspecific body size variation in arthropods over two main macroecological spatial gradients—latitude and elevation—both of high interest in a global warming context. We found that more species with a direct than with an indirect development present a converse Bergmann cline along both gradients. Focusing on spiders, we propose that life history traits such as voltinism, mobility, and brood care influence intraspecific body size patterns—potentially hiding large-scale patterns. Further, we assessed interspecific sexual size dimorphism (SSD) in spider species. We found that extreme SSD—the most original feature in spider biometry—is influenced by hunting guild rather than phylogeny of spider families, suggesting that ecological factors prevail over evolutionary drivers in shaping SSD.

## Introduction

Ectotherms account for approximately 99% of all animal species on Earth, and they represent the vast majority of the living biomass (Atkinson and Sibly 1997, Rosenberg et al. 2023). This diversity and preponderance has to be studied to understand organisms’ ecological and evolutionary responses to environmental changes, such as climate change (Ohlberger 2013). Among ectotherms, arthropods are known for their phenotypic plasticity in life history traits, when habitats vary (Fox and Czesak 2000, Bonte and Lens 2007). As such, they present an ideal taxon to investigate organisms’ responses to environmental changes, which indeed shape morphological and life-history traits expressed in a population (Kingsolver and Pfennig 2007, Nussey et al. 2007). The phenotypic plasticity of arthropods has been demonstrated through variations in life-history traits such as dispersal behavior (Bonte and Lens 2007, Bonte et al. 2008), amount and timing of reproduction events, and clutch size (Fox and Czesak 2000). For example, female spiders have been shown to maximize their fitness by allowing less energy to reproduction whilst producing larger eggs (Hendrickx et al. 2003a, Pétillon et al. 2009). Body size variation has been broadly used to measure the organism responses to environmental variations. It is a relatively easy trait to measure interspecifically and intraspecifically in arthropods because of their exoskeleton, the large number of individuals and species, and their easy collection and preservation. The study of body size, sometimes coupled with geometric morphometrics, gives us information about body condition (Jakob et al. 1996), fecundity (Berger et al. 2012), fitness (Brown et al. 2003), dispersion ability (Coyle et al. 1985) and eating habits (Evans and Forsythe 1985, Gavín-Centol et al. 2017). In the present review we focus on structural body size, as opposed to condition-dependent body size (Box 1). The mean structural body size of individuals from a population is the result of evolutionary and ecological constraints (Gaston and Blackburn 2000), while condition-dependent body size is influenced by the environment of an individual during its whole life. Structural body size (henceforth body size) is a labile trait, i.e. a trait that can vary over the course of an organism’s ontogeny, until the organism finishes its exoskeleton growth (Nussey et al. 2007). Before structural body size is set (i.e before the last ecdysis takes place), it is influenced by physiological traits (growth rate, resources availability, duration of important periods during juvenile growth), themselves influenced by environmental factors (Chown and Gaston 2010) and by phylogeny. Body size variations will be different whether we consider intraspecific or interspercific patterns, generalist or specialist species, and large or small species. It has been shown for example that small ectotherms acclimate to ambient temperature almost instantly (Stevenson 1985). Considering these differences in body size variations, species will be constrained differentially by distinct ecological factors. Body size varies greatly in arthropods: from the 139 µm long male mymarid egg-parasitic wasp *Dicopomorpha echmepterygis* (Mockford 1997, Mymaridae) to the 167 mm long beetle *Titanus giganteus* (Linnaeus 1771, Cerambycidae) (Chown and Gaston 2010). In arthropods, a larger body size can be an asset because it means greater fecundity in females, access to mates for males and resource sequestration advantages (Chown and Gaston 2010, Huang et al. 2017). The benefits of small size may include reduced viability costs of growth and development, enhanced agility and reduced detectability, lowered maintenance energy costs, reduced heat stress, reduced costs of reproduction, and increased scramble competitive ability (Blanckenhorn 2000, Chown and Gaston 2010, Huang et al. 2018). Intraspecific variation of body size and other morphologic traits may ultimately lead to parapatric speciation (Mammola et al. 2018).

The most studied environmental factors in “trait-based ecology of terrestrial arthropods” (Wong et al. 2019) are macroecological factors—namely latitude and elevation—and their associated biotic and abiotic environmental variables, e.g. temperature or season length, often in the context of climate change (Høye and Forchhammer 2008, Radchuk et al. 2013). A latitudinal gradient will trigger gradients of abiotic factors such as temperature and growing season length (Conover and Present 1990) while an elevational gradient will trigger gradients of temperature lapse rate, short-wave radiation input, partial pressure of respiratory gases, precipitation (and thus humidity), turbulence and wind speed (Hodkinson 2005). Biotic factors that vary along these gradients are life history traits such as density, sex ratio (Lee et al. 2012), fecundity (Bowden et al. 2013), dispersal ability, prey availability and sociality (Guevara and Avilés 2007), wing size, color, absorbance and spectral reflectance, body size, thermal tolerance, diapause and fecundity (Hodkinson 2005). Elevational and latitudinal gradients also trigger biotic differences in population density or community composition, themselves having an impact on interactions between organisms and resource acquisition. For example, a higher population density can lead to higher intraspecific competition, leading to a longer development period and a decreased body mass (Begon et al. 1986). Longitudinal gradients are understudied in insects, although they correlate to climatic variables (Jego et al. 2023). Species Abundance and Distribution (SAD) is commonly studied along with body size (Gutiérrez and Menéndez 1997, Gaston 2003, Ulrich 2008, King 2010) and sometimes along with macroecological factors (Ciplak et al. 2008, Bowden et al. 2013). For example, larger repartition area have been shown to positively correlate to body size, showing a direct implication of the study of body size in conservation (Gaston and Blackburn 1996).

Several theories have been proposed to explain the patterns of body size variation along macroecological gradients. The main theory was the Bergmann’s rule which proposes that body size increases with latitude (Atkinson and Sibly 1997), while the converse Bergmann’s rule states that body size should decrease with latitude (Sheridan and Bickford 2011). We present the most studied theories and hypotheses about the role of temperature in Boxes 2 and 3. Along latitudinal gradients, there is a variety of body size responses in arthropods. In a review on 47 species, (Blanckenhorn and Demont 2004) found that, intraspecifically, around 20% of species did not show any pattern, while among the rest, about half followed the Bergmann’s rule, and the other half followed the converse rule. In insects, (Shelomi 2012) found that on a total of 779 intra- and interspecific studies, 30% of the groups followed the Bergmann’s rule while another 30% followed the converse rule and 40% did not follow any trend. If we take some examples, several species of Diptera follow the Bergmann’s rule, while for a lot of Coleoptera, Orthoptera and Lepidoptera (Blanckenhorn and Demont 2004), some spiders (Entling et al. 2010) or univoltine insects (Mousseau 1997), the converse rule has been demonstrated. Elevational gradients have been used as a proxy for latitudinal gradient and climatic changes. Along elevational gradients, body size shows a variety of responses in arthropods, similar to what is observed along latitudinal gradients. Some species show a positive correlation between body size and elevation (Bergmann’s rule) while some others show a negative one (converse Bergmann’s rule) (Hodkinson 2005).

In contrast with arthropods in general, and notably insects, body size variation in spiders is much less known, although they are used in many studies because of their high global and local diversity (see Dimitrov and Hormiga 2021 for a recent review). Like other arthropods, spiders show a very wide size range: the smallest one is thought to be *Patu digua* (Forster and Platnick 1977) from the Symphytognathidae family, and whose body length is 0.37 mm, whilst the largest ones occur among tarantulas (Theraphosidae) and can have body lengths up to 90 mm and leg spans up to 250 mm (Mammola et al. 2017). Gigantism can also be observed on islands in spider taxa such as the *Orsonwelles* genus (Hormiga 2002). As generalist predators, spiders can be used for pest control (Toft 2005). They are good bioindicators (Marc et al. 1999, Rainio and Niemelä 2003) and represent a diverse and abundant group within the arthropods in various habitats (Wise 2006, Lafage and Pétillon 2014). Species with a direct development such as Orthopterans (hemimetabolous insects), and the majority of spiders, have their body size controlled by feeding while ecdysis still takes place; unlike holometabolous insects (with indirect development) such as ground beetles whose imaginal body size is governed only by larval stages (complete metamorphosis). Spiders body size and shape have been shown to vary across behavioral guilds (Wolff et al. 2022). Along macroecological gradients, we hypothesize that wanderers and web-building species might present different patterns of body size variation. In spiders, males are typically smaller than females and present one of the greatest sexual size dimorphisms (SSD) in animals (Kuntner and Coddington 2020) (Figure 1). Only in rare cases is the male larger than the female (Aisenberg et al. 2007). Across the animal kingdom, SSD tends to increase with mean body size when the male is the larger sex; while the opposite tends to arise when the female is the larger sex. This is known as the Rensch’s rule (Rensch 1950, Bidau and Martí 2008) and has also been studied intraspecifically and along latitudinal gradients (Blanckenhorn et al. 2006, Høye and Hammel 2010, Bowden et al. 2013, Laiolo et al. 2013). While most arthropod species follow the Rensch’s rule—with some notable exceptions (Bidau et al. 2016, García-Navas et al. 2017, Rossi and Haga 2019)—spiders do not follow it (see e.g. Figure 1c in Kuntner and Coddington 2020). Several hypotheses have been advocated to explain extreme SSD in spiders (summarized in Foellmer & Moya-Laraño, (2007)). Selection may favor smaller males because (i) being mature earlier than the females (protandry) is advantageous, (ii) larger specimens are eaten by the females (cannibalism) (although this hypothesis has been refuted multiple times, see Foellmer & Moya-Laraño, (2007)), and (iii) a smaller body size allows males to more easily climb to high habitat patches where females are located (gravity hypothesis) and thus win the scramble competition, the race for reaching mates by the time they are receptive (Moya-Laraño et al. 2002). Selection may in parallel favor larger females because (i) they have higher fitness, (ii) they get predated less (predation release (Hormiga et al. 2000, Higgins 2002)), and (iii) they profit from larger prey availability (Foellmer and Moya-Laraño 2007). Overall, extreme SSD in spiders is likely due to divergent selective directions on males and females (Kuntner and Coddington 2020). As in spiders, extreme female-biased SSD is found in mantis, where it has been shown that female body size was related to prey size, fecundity, and emergence, and not related to cannibalism or male attraction (Maxwell and Frinchaboy 2014). Yet, most studies assessing processes driving SSD considered single spider species, or reviewed species in local parts of the world (Foellmer and Moya-Laraño 2007).

**Figure 1.**
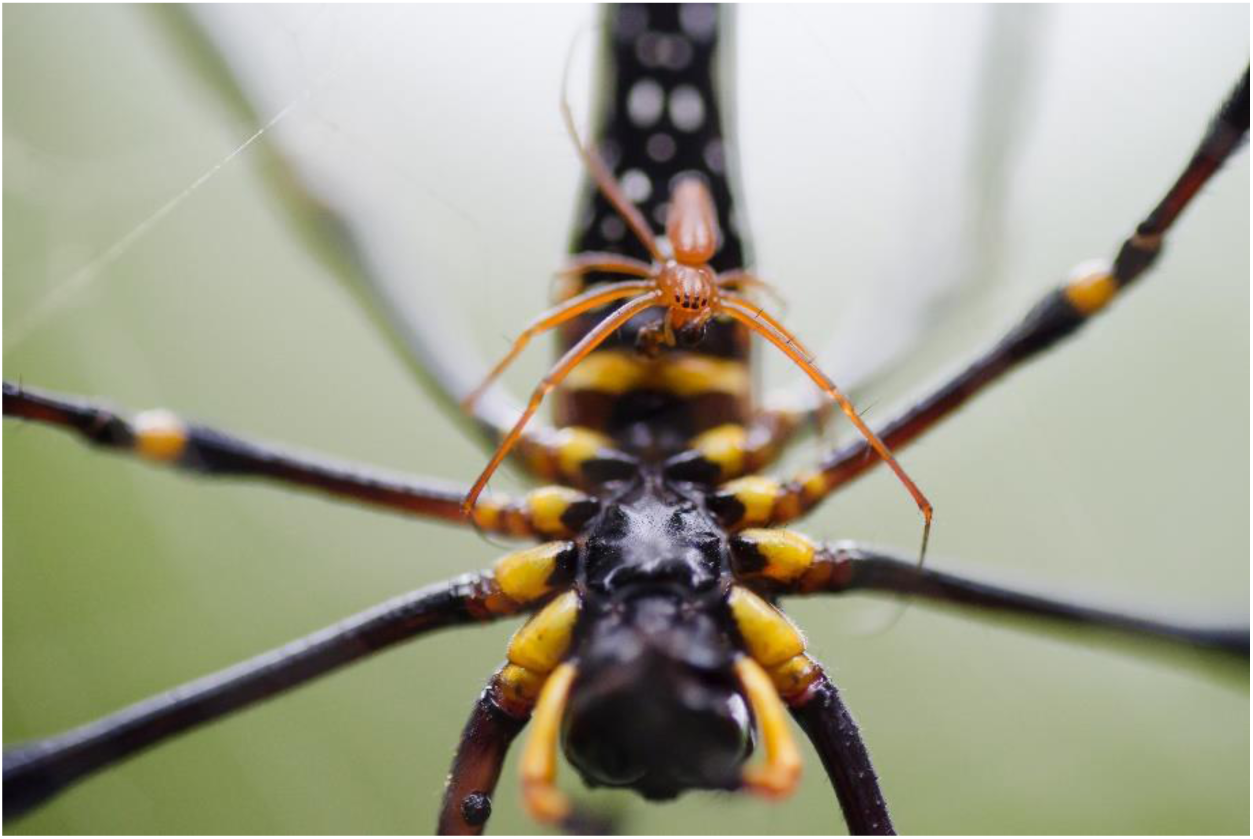
Sexual size dimorphism (SSD) in spiders. A male (above) and a female (below) Golden Orb-web spider *Nephila pilipes* mating in the Kaeng Krachan National Park, Thailand. Photograph by Thai National Parks, distributed under a Creative Commons Attribution 2.0 Generic license.

In this work, we review field studies on intraspecific variation of structural body size along macroecological gradients in spiders and other arthropods. We extend the analysis from (Foellmer and Moya-Laraño 2007)—using a large database of 1822 worldwide species synthetizing spider traits and phylogeny—to test hypotheses on both ecological and evolutionary drivers of spider SSD. We address the following three questions: (1) How do body size and SSD of spiders vary along macroecological factors? (2) Is this variation different from other arthropods and among taxa with distinct development (direct vs. indirect)? (3) How do biotic (including life history traits) and abiotic factors interact with body size and SSD, to possibly explain larger spatial scale patterns? We further discuss the implications of monitoring spider body size and SSD in a changing world.

## Methods

### (1) Macroecological patterns in arthropods and spiders

We searched for reviews and articles using the search engine ‘Google Scholar’ until September 2024 with the following key words: ‘body size’ + ‘macroecological’ / ‘altitude’/ ‘latitude’ / ‘elevation’ + ‘ectotherm’ / ‘Arthropod’ / ‘insect’ / ‘spider’/ ‘Hexapoda’ / ‘Chelicerata’ / ‘Arachnid’ / ‘Araneae’ / ‘Myriapoda’ / ‘Crustacea’. This resulted in 40 combinations of searches. Specifically for spiders, we report studies that we found on other microecological patterns (urbanization and habitat quality) as a non-exhaustive list.

### (2) Sexual size dimorphism

Data on species mean body size were extracted from the betsi database (https://portail.betsi.cnrs.fr/). Data on species substrate were extracted from the Spider and Harvestman Recording Scheme website (http://srs.britishspiders.org.uk/). To avoid a possible size effect due to varying species richness in families we only used families with at least ten species, resulting in a total of 48 families (432 genus, 1822 species). Each family was attributed a guild, either web-builders or non web-builders, based on (Cardoso et al. 2011). Correlation between female body size and female/male body size ratio was estimated in a Bayesian framework with a Student’s t distribution with the brms package (Bürkner 2018). We used 2000 iterations on four chains. Model convergence was checked by visually inspecting diagnostic plots. The link between female body size and female/male body size ratio was estimated using models built within a Bayesian framework using brms (Bürkner 2018) with two chains and default priors (except for intercept which was set as normal (1,1)). The models included female body size only, female body size and family, female body size, family and guild, guild and size as predictors. Family was included as a random factor. Model convergence was checked by visually inspecting diagnostic plots and using Rhat value. Parameter selection was based on “HDI+ROPE decision rule” with a range value determined as -0.1 * sd(y), 0.1 * sd(y) (Kruschke and Liddell 2018) and was performed using bayestestR (Makowski et al. 2019). The best model was selected based on LOO comparison.

## Definitions and concepts

### Box 1.

Body size

It is important to differentiate structural body size (SBS, usually called body size indicator (BSI) in vertebrates) from condition-dependent body size (CDBS). Structural body size was defined by (Moya-Laraño et al. 2008a) as a ‘*measure of body size that does not change with additional nutrient acquisition once an animal has discontinued growth, and thus SBS is not directly affected by current condition*’, while CDBS is ‘*any measure of body size that changes with current condition (i.e. covaries with nutrient storage)*’. Body condition (i.e. body mass corrected for size differences) is a proxy of fitness and reflects nutrient storage and habitat quality (Jakob et al. 1996, Moya-Laraño et al. 2008a, Peig and Green 2009, Knapp and Knappová 2013, Labocha et al. 2014). Structural body size, on the other hand, is not affected by condition and will thus reflect evolutionary and ecological constraints. In arthropods, the confusion is sometimes made between CDBS and SBS when body condition traits are measured and considered as ‘body size’ such as mass, density, or abdomen size (Chown and Gaston 2010).

Structural body size can be measured in multiple ways within and across taxa, considering distinct ways to store energy. For example, the whole body of a ground beetle is covered with chitin, which prevents it from storing much energy while a spider’s abdomen can grow larger to store energy. It will then be more adapted to measure a sclerotized body part such as the head or the pronotum to measure SBS. Measures of structural body size in spiders include: cephalothorax (carapace) length (Petersen 1950) and width (Høye and Hammel 2010, Bowden et al. 2013) illustrated in Figure 2A: *Pardosa furcifera*, from Ameline *et al*., (2017), first femur length (Opell et al. 2007) (illustrated in Figure 2B), and tibia-patella length (Skow and Jakob 2003). There needs to be a consensus on which body size measures to use in order to allow comparison (Gallagher et al. 2020). Indeed, interspecific meta-analyses using average figures from taxonomic literature (Ulrich 2007, Entling et al. 2010, Ulrich and Fiera 2010, Fattorini et al. 2013, 2014) can only do so if the same measures are used. Similarly, intraspecific studies should be comparable among one another. In most recent studies, several measures of sclerotized body parts are correlated (Knapp and Knappová 2013, Puzin et al. 2014, Ameline et al. 2018, Lowe et al. 2020). In the present study, we are interested in evolutionary and ecological constraints, thus focusing on structural body size, henceforth body size throughout this article.

**Figure 2.**
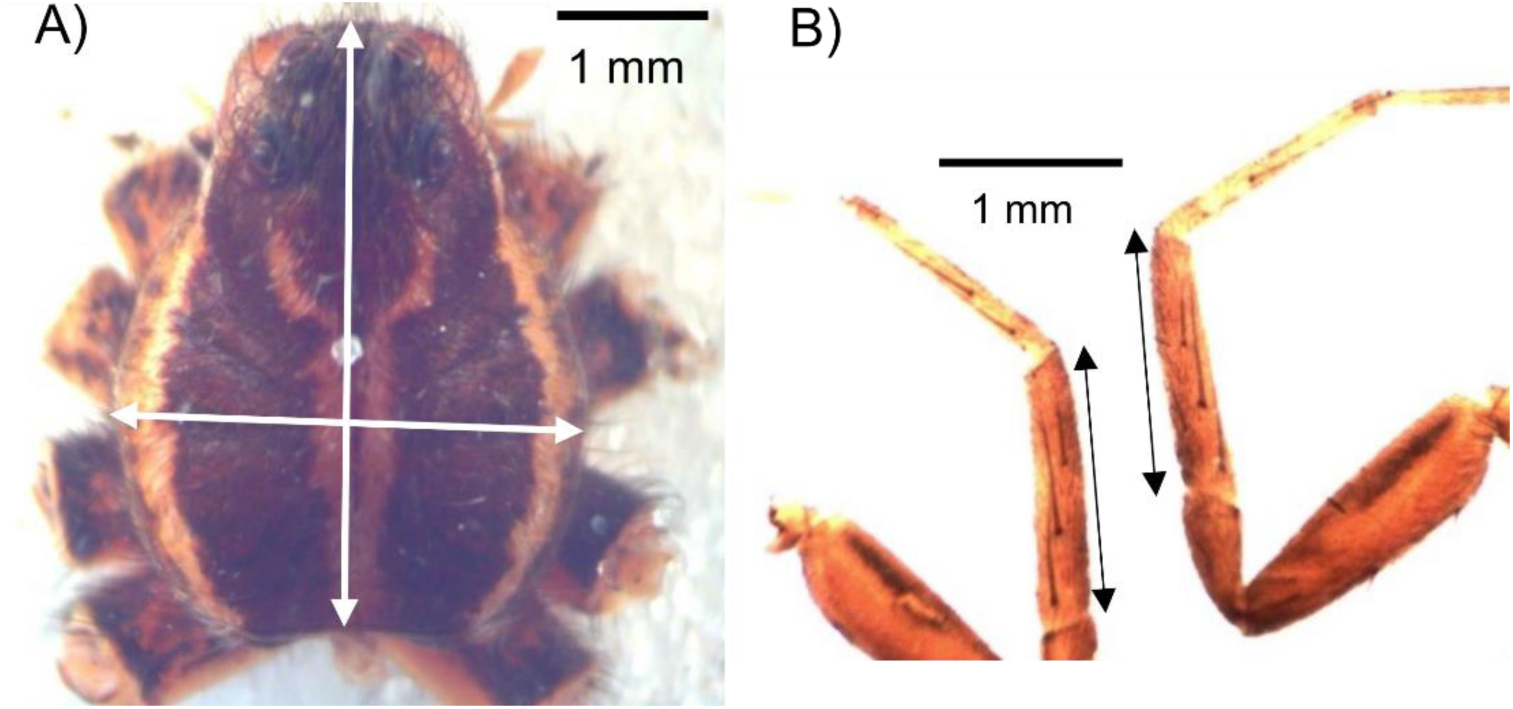
Structural body size measurement in the wolf spider *Pardosa furcifera* (Araneae, Lycosidae). From Ameline *et al*., (2017). **A**: carapace length and width, **B**: first femur length.

### Box 2.

Rules of body size variation along macroecological gradients

Body size has been used interspecifically in allometric studies to try and understand general biological laws (Norris 1998, Raerinne 2013). At the interspecific level, the *Metabolic Theory of Ecology* proposes that (i) there are more species at low latitudes than at high latitudes, (ii) there are more small species, (iii) so there are consequently more small species at low latitudes and they show greater diversity (Brown et al. 2004). The most famous body size variation theory is the *Bergmann’s rule* which proposes that body size increases with latitude (Bergmann 1848, Atkinson and Sibly 1997). This rule states that a larger organism provides a reduced surface per volume ratio which minimizes the heat loss (Blackburn et al. 1999). The rule was originally thought to apply interspecifically to endotherms, but an intraspecific version has been proposed as *James’ rule* or *neo-Bergmann’s rule* (Chown and Gaston 2010, Vinarski 2014), and it has been tested in all taxa (Blanckenhorn and Demont 2004, Chown and Gaston 2010, Entling et al. 2010, Meiri 2011, Shelomi 2012). As the opposite pattern has been observed in arthropods (Sheridan and Bickford 2011) and spiders (Puzin et al. 2014), a *converse Bergmann’s rule* has then been proposed which postulates that decreasing season length causes smaller body size at higher latitude (Blanckenhorn and Demont 2004). This rule supposes that shorter seasons at higher latitudes reduce the foraging period, hence resulting in reduced growth and development period. A gradient of decreasing body size towards the poles should then be observed (Blanckenhorn and Demont 2004). The *countergradient variation* announces that growth rate increases with latitude, because the growing season is shorter, resulting in a homogenous distribution of body size along latitudinal gradients (Conover and Present 1990). Different intraspecific patterns of body size variation across macroecological gradients are usually reported, including linear and non-linear clines. Linear clines can be *positive* (Bergmann’s rule) or *negative* (converse Bergmann’s rule) showing increase or decrease of body size across the range, respectively. Non-linear clines include *sawtooth* and *stepwise* clines (Figure 3A and B, from Shelomi, (2012), which can still show an overall positive or negative cline, *U-shaped* or *hump-shaped* clines, which occur when an increase or decrease is reversed gradually over a single point in the entire range (Figure 3C and D), while *curvilinear* clines occur when increase or decrease is reversed gradually over several points across the range (Shelomi 2012). The *Rapoport’s rule* says that the size of a species distribution range increases with latitude, but it is used interspecifically (Gaston and Blackburn 1996). The *Cope’s rule* supposes a macroevolutionary trend towards increased size (Kingsolver and Pfennig 2004). The *seasonality hypothesis* states that climates characterized by long annual cycles select for large body sizes. This hypothesis was confirmed by (Troost et al. 2009) who proposed that both longer productivity and seasonality may lead to larger body sizes. Although many hypotheses were proposed, the mechanisms underlying these body size patterns are still rarely investigated (Blanckenhorn and Demont 2004).

**Figure 3.**
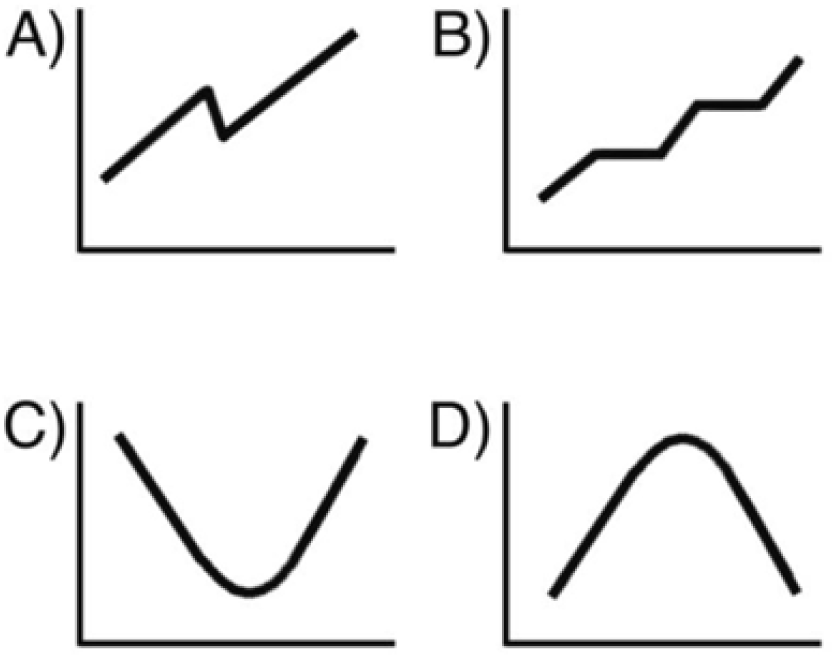
Non-linear clines as described in (Shelomi 2012). Non-linear clines include *sawtooth* and *stepwise* clines (**A and B**), which can still show an overall positive or negative cline, *U-shaped* or *hump-shaped* clines, which occur when an increase or decrease is reversed gradually over a single point in the entire range (**C and D**), while *curvilinear* clines occur when increase or decrease is reversed gradually over several points across the range.

### Box 3.

Body size variation linked to temperature variation

Rules laid down are often thought to be linked to temperature. However, it is still a controversial subject and there is no clear explanation on why body size increases or decreases when the organism is exposed to low temperatures (Atkinson and Sibly 1997). Indeed, 83% of 109 studies on ectotherms showed an increased growth rate and consequently, an increased body size at lower rearing temperatures, although individual growth rates were expected to decrease at lower temperatures. This phenomenon is explained by the *‘developmental temperature-size rule’* or *TSR* (Atkinson 1994, Atkinson and Sibly 1997). The organism can ‘choose’ to trade-off for a delayed maturity which would lead to a larger body size if the increase of fecundity exceeds the loss of fitness caused by delayed maturity. Another hypothesis is that higher temperatures may cause limited resource availability. In response to that limitation, body size is smaller due to the lack of resource acquisition or due to an adaptive response to reduce size at maturity. From a cellular point of view, the TSR argues that cellular differentiation which leads to sexual maturity is limited by diffusion at low temperatures, thus maturity would be delayed. Consequently, cell size would become larger at lower temperatures (Arendt 2007). The Bergmann’s rule has been attributed to the TSR theory (temperature variation) while the converse Bergmann’s rule was attributed to a resource limitation, due to season length variation (Hodkinson 2005). This has been shown, for example, by an experiment along a thousand meters elevational gradient on weevils in subantarctic islands by (Chown and Klok 2003).

Kingsolver and Huey (2008) summarized the rules linked to temperature as three rules. The first one is “*bigger is better*” (red in Figure 4, adapted from Kingsolver & Huey, (2008)). A larger body size is thought to bring a higher fitness via better performance and dominance. But a larger body size can also be costly and risky because of delayed maturation and increased energy demand. Hence, intermediate sizes are also thought to be optimal. As a counter-example of this first rule, (Moya-Laraño et al. 2007) showed that smaller males are better scramble competitors. Smaller is also generally better in the case of parasites and parasitoids (Blanckenhorn 2000). The second rule is “*hotter is smaller”* (green in Figure 4). It corresponds to the TSR, which takes in account the effect of temperature on the organism during development. Counter-examples exist, e.g. in the case of Orthoptera and Lepidoptera, hotter is bigger (Mousseau 1997, Walters and Hassall 2006). ‘*Ectotherms that obey the temperature size rule are identified as having a higher temperature threshold for development rate than for growth rate; exceptions are identified as having a lower temperature threshold for development rate than for growth rate*’ (Walters and Hassall 2006). The third rule, “*hotter is better*” (blue in Figure 4), proposes that organisms with a high optimal temperature will also have high maximal performance and fitness, according to thermodynamics laws. We chose not to present body size variations in response to temperature because most of the studies consider growth rate and growth efficiency but not directly body size (Angilletta and Dunham 2003).

**Figure 4.**
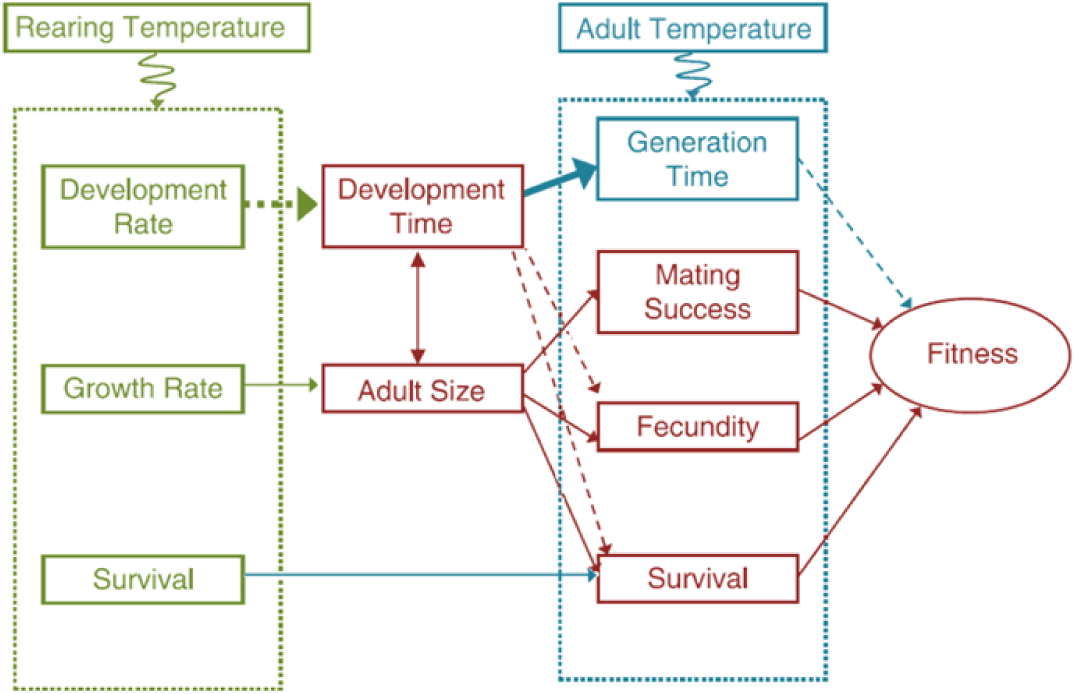
Body size variation linked to temperature variation. Adapted from Kingsolver & Huey, (2008). Red: “*bigger is better*”, green: “*hotter is smaller*”, blue: “*hotter is better*”.

## Results and discussion

### (1) Body size and sexual size dimorphism (SSD) along macroecological gradients

We found a total of 146 studies reporting intraspecific body size variation of 240 arthropod species along latitudinal and elevational gradients (Figure 5, Supplementary Table S1). We did not find an overall tendency towards positive (Bergmann’s rule) or negative (converse Bergmann’s rule) variation of body size across elevation or latitude. We found that species with a direct development presented a larger proportion of negative clines than species with an indirect development, both along elevational (Z-test, X-squared = 4.1, df = 1, p-value = 0.02, 42% vs. 23%, Figure 5) and latitudinal clines (Z-test, X-squared = 8.2, df = 1, p-value = 0.002, 74% vs. 40%, Figure 5). This result suggests that the growth of species with a direct development might be restricted by the environmental changes caused by increasing latitude or elevation—notably the shorter season length. Because species with an indirect development go through metamorphosis between the larval and the adult stages, they may dispose from more possibilities to increase their growing season. In spiders, we found that 11 out of 18 species showed no body size trend along elevational gradients (61 %, Figure 5). This proportion was higher than in other arthropod species (Z-test, X-squared = 8.2, df = 2, p-value = 0.02, 61% vs. 22% and 33% in species with direct and indirect development, respectively). This result may be caused by the high dispersal of spiders which would hide existing body size variations along elevational gradients. It is important to note that most studies have been performed on a limited number of populations and have not been repeated, which may result in a low statistical power to detect overall patterns (Gienger et al. 2019).

**Figure 5.**
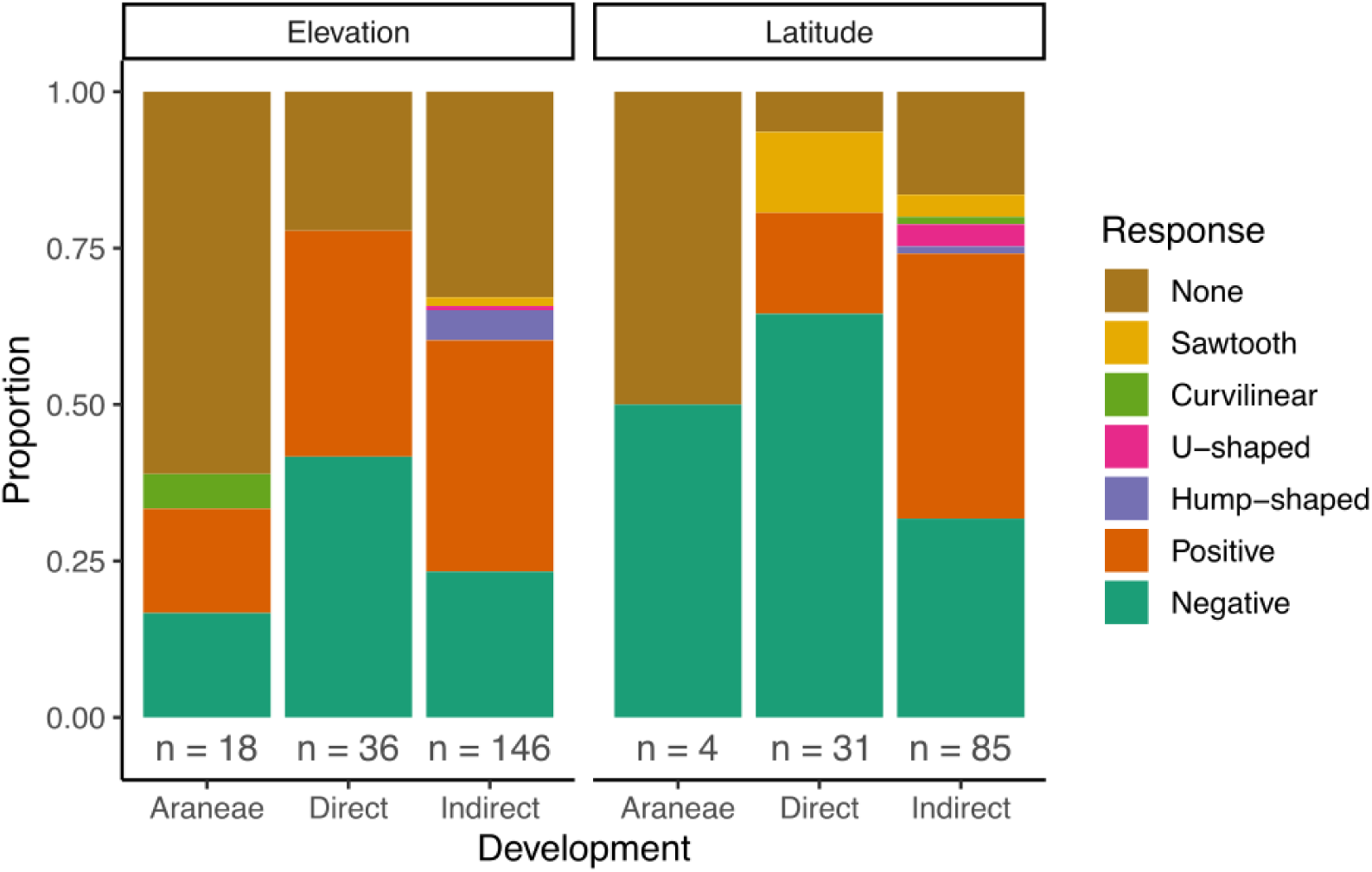
Intraspecific body size response to elevation and latitude in arthropods. We separately report studies on spider species along elevational gradients. Only two studies were found in spiders along latitudinal gradients, we included them in the direct development category. We distinguished taxa with direct from taxa with indirect development (Schäfer 1976).The number of species is reported below each barplot. Details of the studies are given in Supplementary Table 1.

Sexual Size Dimorphism (SSD) has been shown to vary across macroecological gradients in arthropods interspecifically (Blanckenhorn et al. 2006, Laiolo et al. 2013) and intraspecifically, e.g. in beetles (Baranovská et al. 2019) and spiders (Høye et al. 2009, Høye and Hammel 2010, Bowden et al. 2013, Laiolo et al. 2013). A meta-analysis on SSD variation across macroecological gradients in arthropods revealed that, on average, males body size varied more than females (Horne et al. 2019, Kojima 2019). In spider species this was found to be the opposite, as female body size varied more across elevation gradients (Horne et al. 2019). This study suggests that SSD variation is constrained by evolutionary forces rather than environmental ones (Horne et al. 2019). They also found no relationship between SSD variation and varied tested factors like environment type (aquatic or terrestrial), voltinism, mean body size, degree of SSD and gradient direction. Conversely, it was found in grasshoppers that female body size varied more across an elevational gradient and the authors hypothesized that female body size might be more plastic or evolve faster in harsh environments (Laiolo et al. 2013).

In contrast with sexual *size* dimorphism, sexual *shape* dimorphism is studied in other arthropods (Vujić et al. 2020) and might be relevant in further, spider-focused, studies.

In the context of anthropic impacts, macroecological gradients have been studied in spiders, such as urbanization and land use. It was shown for one species of orb-weaver spider (Araneidae) that body size increased in an urbanized habitat (Lowe et al. 2014), while another species was shown to decrease body size in highly urbanized habitats (Dahirel et al. 2019). Interspecifically, urbanization and fragmentation have been shown to cause body size decrease (Merckx et al. 2018, Piano et al. 2020). High habitat quality, determined by more rocks, precipitation and duration of snow coverage, was shown to intraspecifically increase body size and eggcase size (Mammola et al. 2019). Corroborating this result, (Bowden et al. 2018) observed a decline in spider abundance correlated to snowmelt timing and moisture availability. In an agricultural context, (Plath et al. 2021) revealed that mean body size of spider community decreased with intensive agriculture. On the other hand, (Pinto et al. 2021) observed no difference in body size in spider communities across different land uses, and hypothesized that microhabitat rather than global land use influenced the observed interspecific body size pattern. In the following, we pinpoint several factors that could explain variability of intraspecific body size responses along macroecological gradients among taxa and even species, focusing on spiders.

### (2) Biotic and abiotic factors driving local changes in body size and SSD

#### (a) Biotic factors

Diet quality has been correlated to body size variation across latitudinal gradients (Ho et al. 2010). Higher competition and variation in prey availability might also lead to fewer resources and explain a lack of overall body size variation along environmental gradients (Beckers et al. 2020).

Among life history traits, the degree of voltinism—i.e. how many reproduction events occur during the year—may influence body size (of both male and female) response to macroecological gradients. Shifts in voltinism across elevational or latitudinal gradients—due to changing seasonality—might cause body size variation along these gradients (Beckers et al. 2020). (Cabon et al. 2024) showed that temperature had an effect on the body size of univoltine, but not bivoltine species. A sawtooth cline is typically interpreted as the result of a shift in voltinism (Shelomi 2012). The level of voltinism is known to change interspecifically with latitude and hence temperature (Kivelä et al. 2011, Zeuss et al. 2017). Voltinism is less studied across populations and along elevational gradients, as these gradients are usually smaller than latitudinal ones. A shorter generation time in multivoltine species would provide a longer growing season for earlier clutches of offspring (Chown and Klok 2003). A different degree of voltinism would then mean a different resource allocation strategy and the underlying mechanisms of body size variation would differ. In the present study, we could not consider voltinism *a priori*, as it may vary within species and populations.

Brood care is common in spiders and may provide them with a lower susceptibility to unfavorable conditions in comparison with insect species with stationary larval stages (Beckers et al. 2020). Body size might then be less influenced by environmental conditions during development and show no trend across an environmental gradient.

Mobility and dispersal are important factors that could influence the degree of body size variation across gradients. In (Cabon et al. 2024), only species with a low ballooning capacity showed body size variation across a temperature gradient. Species with low mobility may be split in subpopulations that would induce body size variation across gradients (Beckers et al. 2020). A higher mobility may thus cause a species to show variation induced by large-scale rather than fine-scale environmental factors (Hein et al. 2019). For example, body size variation has been linked to foraging ability in bees (Gathmann and Tscharntke 2002). Here we want to emphasize the role of body size particularly in spider dispersion. Long-distance dispersal in spiders will be passive via ballooning and favored by a small body size or mass while short-distance dispersal will be active via walking on the ground and favored by a large size or amplitude. Ideal Free Distribution (IFD) is expected to homogenize body sizes among habitats (Fretwell and Lucas 1970, Hendrickx et al. 2003b, Yip et al. 2008), while disturbances may select for increased dispersal propensities: smaller body size in ballooning species or individuals, mostly web-building spiders (Coyle et al. 1985, Blandenier 2009, Blandenier et al. 2013, Simonneau et al. 2016) or longer legs in runners, i.e. ground-dwelling spiders (Lafage et al. 2015). Selection for relatively longer legs was previously reported in web-building (mate search: Foellmer and Fairbairn 2005), free-running (mate search: Framenau 2005) and cave-living species (cave adaptation: Miller 2005). Longer legs have also been proved to increase speed in steep ground (Prenter et al. 2012), and in suspensory locomotion (Moya-Laraño et al. 2008b). Although we do not present an exhaustive list of such results, traits linked to body size have been shown to vary across macroecological gradients: sexual size dimorphism (SSD), growth rate—related to temperature in (Angilletta, Jr., and Dunham 2003), or fecundity (Reed and Nicholas 2008, Berger et al. 2012, Bowden et al. 2013, Marshall et al. 2013).

Body size in spider offspring is typically measured to reveal reproduction trade-offs (Petersen 1950, Brown et al. 2003, Skow and Jakob 2003, Walker et al. 2003, Pétillon et al. 2009, Ameline et al. 2017), and these reproductive trade-offs have been revealed along macroecological gradients (elevation: Ameline et al. 2018).

#### (b) Abiotic factors

Habitat has been shown to be an important factor in spider communities distribution (Koponen 2002, Entling et al. 2007). At a local spatial scale (m^2^), habitats can shape body size patterns (Brown et al. 2003, Hendrickx et al. 2003a, Pétillon et al. 2009, Puzin et al. 2011). After finding no overall body size patterns along elevational gradients, studies suggested that microhabitats played a more important role than the large-scale landscape (Hein et al. 2015, Ameline et al. 2017). Local environmental conditions such as soil pH, occurrence of ice and abundance have been shown to play a role in species body size distribution (Ornaghi et al. 2023). In spiders, snow cover and habitat type are considered to play a role in body size distribution across macrogeographic scales (Hein et al. 2014, Bowden et al. 2015). (Bowden et al. 2015) emphasize that environmental conditions are especially important for small size species with low dispersal ability. Further, models showed that body size was linked to how ectotherms move in microclimatic environments (Woods et al. 2015).

Climate has been shown to drive body size patterns in arthropods along macroecological gradients (Hu et al. 2012). Alternatively, little effect of climate change on body size has been observed in herbivorous beetles (Baar et al. 2018). The effect of temperature on body size or related traits is often studied in insects (Opell et al. 2007, Ciplak et al. 2008) and has also been studied along latitudinal and elevational gradients (Koyama et al. 2015, Pérez-Valencia and Moya-Raygoza 2015). In insects, further analysis of the effect of temperature on physiology (growth rate, metabolic rate) has been performed along elevational gradients (Bulgarella et al. 2015, Cogo et al. 2020). Dispersal ability, mentioned above, has also been shown to be influenced by temperature in spiders (Bonte et al. 2008). In various arthropods, temperature and season length has been found to influence body size—e.g. in crustaceans (Jaramillo et al. 2017) and beetles (Collard et al. 2021). Overall patterns of reduced body size were observed with warming temperatures, likely linked to oxygen lack in aquatic environments (Verberk et al. 2021) and heat tolerance in endotherms (Peralta-Maraver and Rezende 2021).

Other climatic factors such as photoperiod (Hassall 2013) have been correlated to body size variation along macroecological gradients. We further develop the broader impact of climatic changes on body size variation in the last section of the results.

In this review, we collected data about interspecific SSD in spiders, using a large dataset of 48 families (432 genus, 1822 species). SSD was previously shown to be positively correlated with female size in spiders (converse Rensch’s rule: Foellmer and Moya-Laraño 2007). Using a larger dataset, we confirm this hypothesis (Figure 6). Across spiders species, body size has been shown to be highly phylogenetically conserved (Entling et al. 2007). We thus use mean body size of males and females in spider species to approximate interspecific SSD.

**Figure 6.**
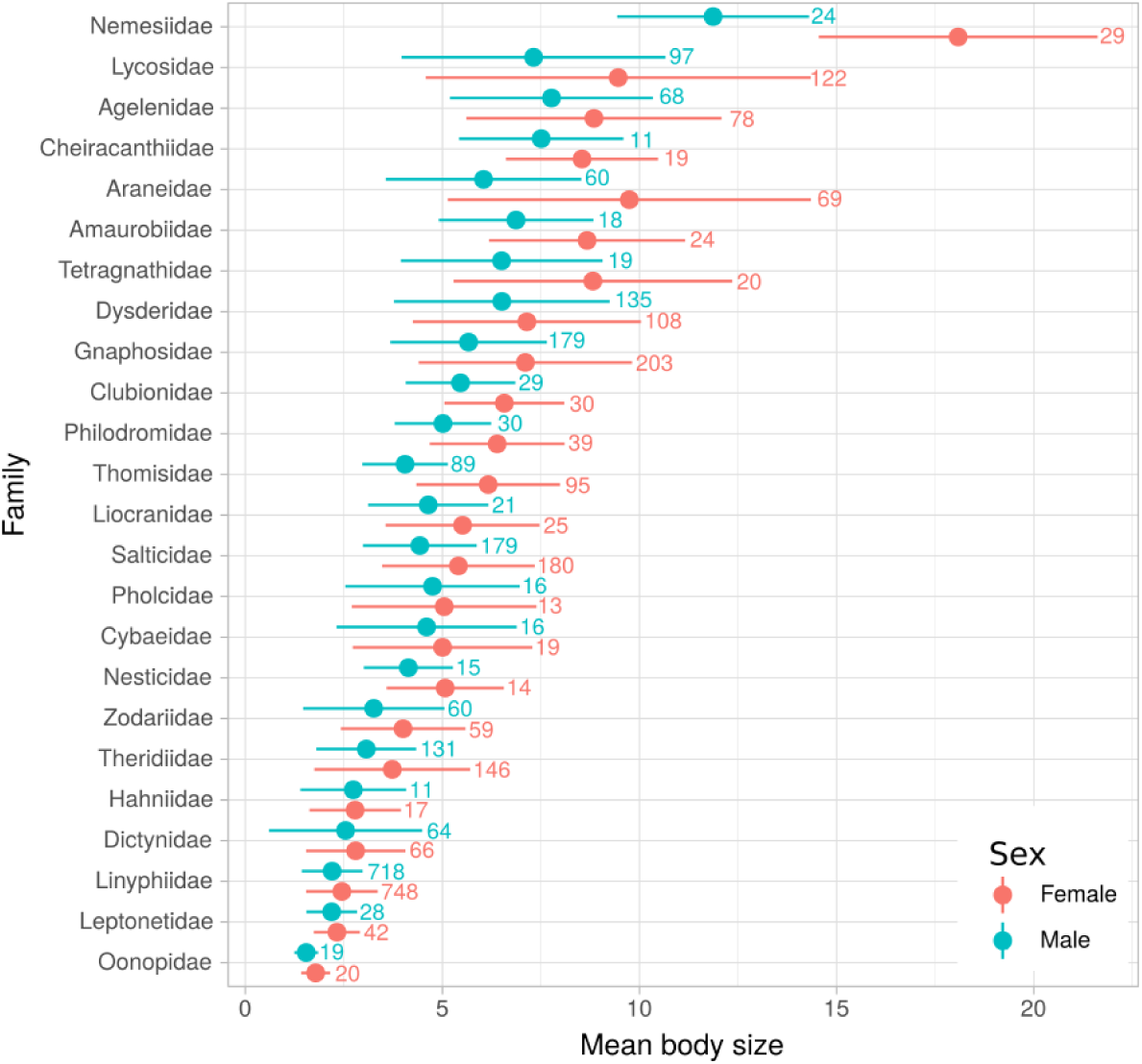
Correlation between sexual size dimorphism (SSD) and mean body size in spider species.

The ratio between female and male body size was significantly correlated with female body size (R² = 0.42, 95%-CI = 0.38 – 0.46). The best model explaining the ratio between female and male body size included female size and family as predictor with a R² of 0.327 (95%-CI = 0.289 – 0.357). Female size effect on the ratio between female and male body size has a very high probability of existing and being positive (pd = 100%, Median = 0.032, 89% CI [0.028, 0.035]) and could be considered as significant (0% in ROPE). Family effect on the ratio between female and male body size has a very high probability of existing and being positive (pd = 100%, Median = 0.21, 89% CI [0.20, 0.22]) and could be considered as significant (0% in ROPE). We found that SSD was less prominent in web-building spider families (Figure 7, Mann-Whitney test, W = 554928, p < 0.0001). We found no curvilinear relationship between ln (males) and ln (females) in either web-building and non web-building spiders (Figure S1) contrary to the expectations regarding the ‘gravity hypothesis’ made by Foellmer & Moya-Laraño (2007). We found no correlation between SSD and more specific guild categories within web-building and non-web building species (Figure S2, Mixed linear model: ratio∼guild+1|family, X^2^ = 16.13, df = 7, p = 0.024). We then tested whether the position of the species in the substrate (ground, herbaceous, shrub, canopy) was correlated with SSD (Figure 8) and found no correlation (Mixed linear model: ratio∼substrate+1|family, X^2^ = 13.44, df = 7, p = 0.062). To test the correlation between SSD and the phylogeny of spider families, we projected the mean values of SSD on a phylogenetic tree (based on Macías-Hernández *et al*., (2020)) for each family containing more than ten species (Figure 9). We did not find that phylogeny influenced SSD in spider families. Our results suggest that ecological rather than evolutionary factors influence SSD in spider species.

**Figure 7.**
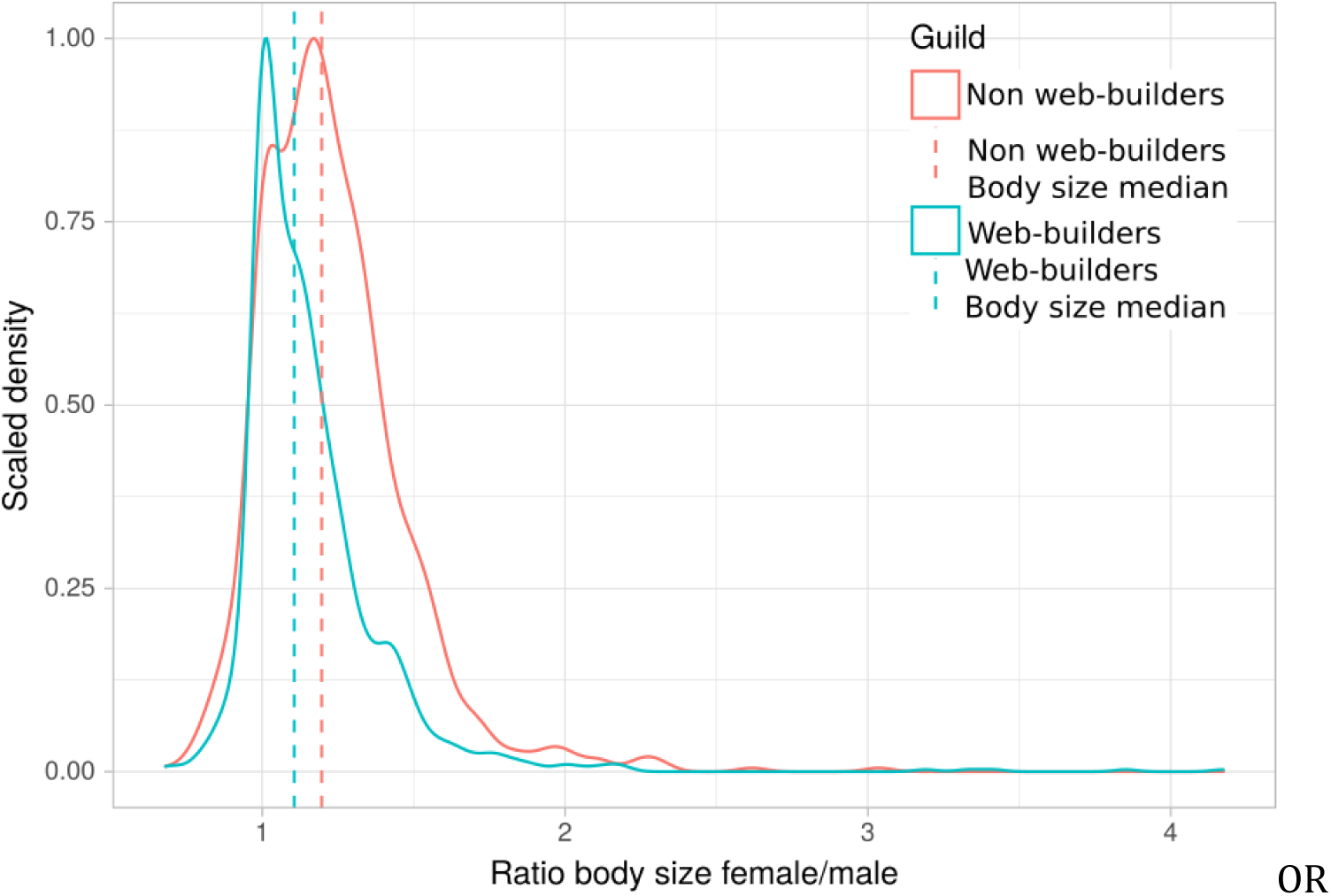
Sexual size dimorphism in non web-building and in web-building spiders.

**Figure 8.**
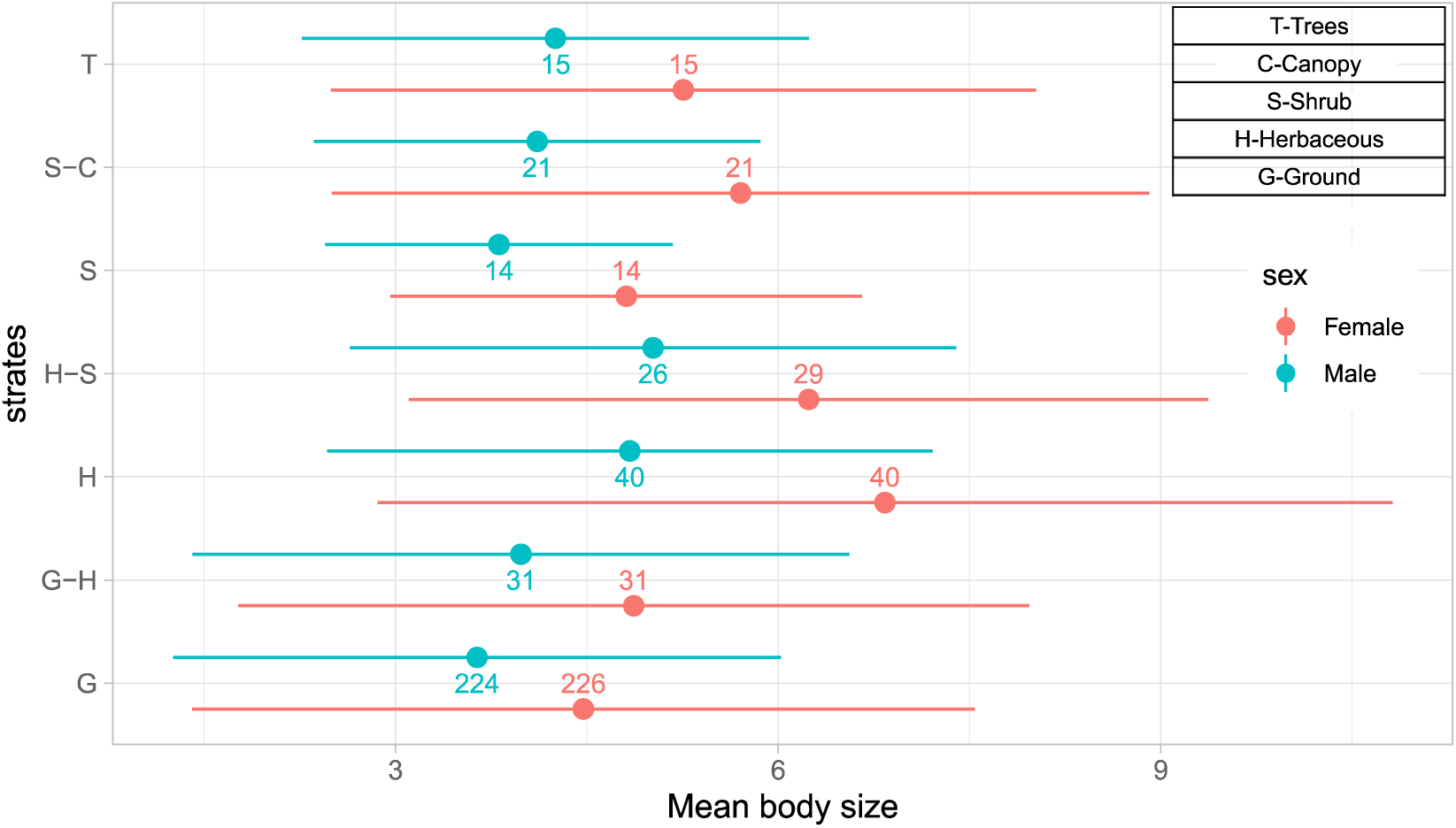
Sexual size dimorphism in spider species occupying distinct substrates.

**Figure 9.**
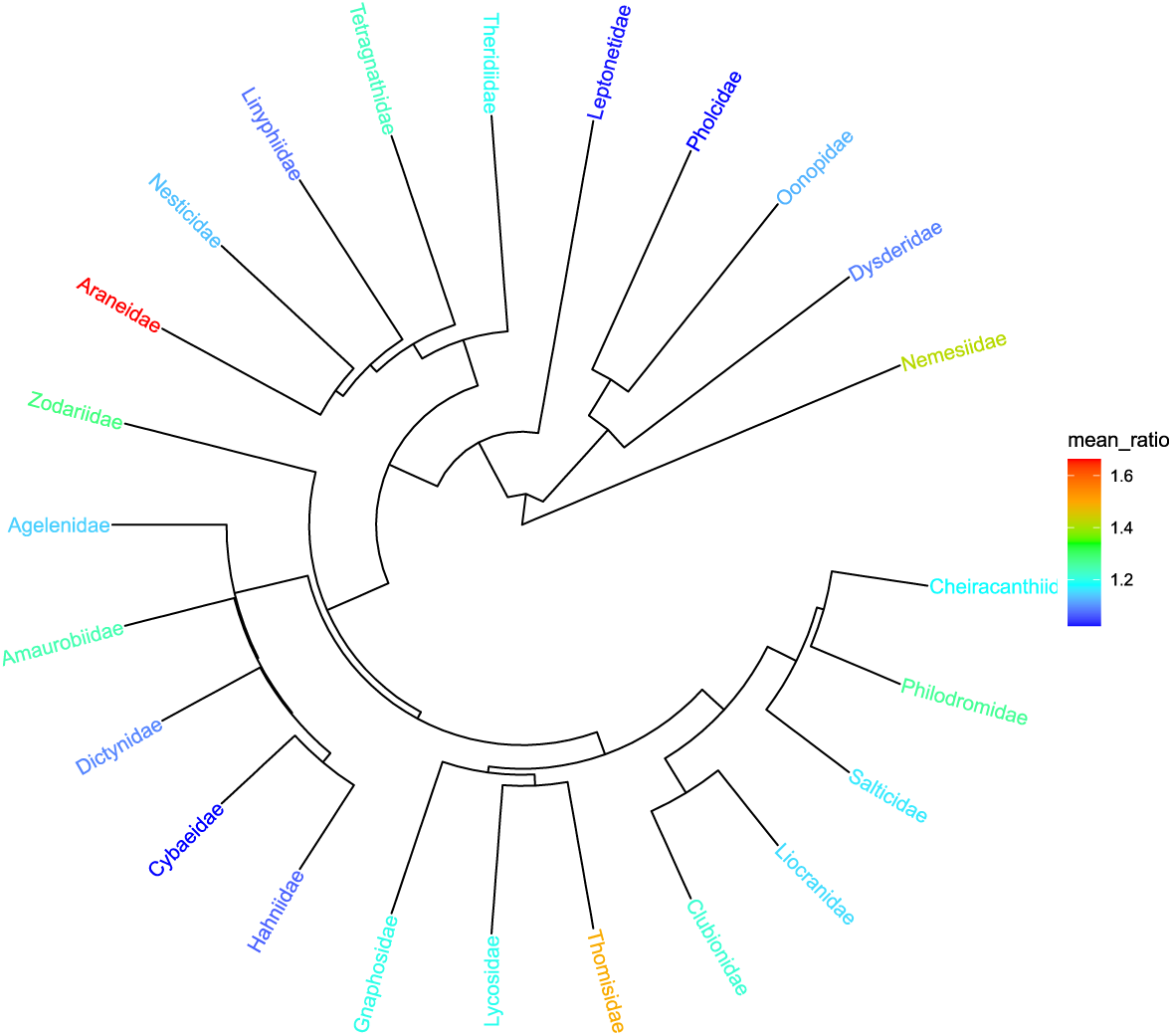
Sexual size dimorphism projected on the phylogenic tree of spider families.

### (3) Perspectives and future research directions in a changing world

Latitude and elevation are macroecological factors that are good correlates of temperature variations in a global change context. There is a need to predict how species and populations will adapt to macroecological variations, since global change brings fast and important modifications. Anthropic activity threatens biodiversity and thus stability of ecosystems and biogeological cycles, causing an increase of severe weather events. It has been shown for birds that severe weather events can exert directional selection pressure, favoring larger size (Goodman et al. 2012). Global warming is thought to generally result in an intraspecific decrease in body size, but the underlying mechanisms are not well known and very few studies have focused on endotherms (Sheridan and Bickford 2011). Body size and SSD can also vary with anthropogenic disturbance (Sukhodolskaya and Eremeeva 2013). Intraspecifically, climate change may result in increased SSD at higher elevations due to expanding growing seasons (Høye and Hammel 2010).

Thermal tolerance may be studied along with body size variation (Hoffmann et al. 2013), although we assume that it may play a less important role than habitat niches in ectotherms. Sudden environmental changes can lead to niche shifts, thus leading to invasions depending on dispersion ability and range size (Iversen et al. 2013). Climate indeed acts as a major constraint on range boundaries (Thomas 2010). This is a major social issue as invasive species may also be vectors of disease, such as mosquitoes with malaria. Invasions can also inverse the clines brought by climate change (Treasure and Chown 2014). A good example of habitat that is particularly prone to global warming is the arctic habitat; where arthropods rely on snowmelt to emerge. Hence, the predicted variability of snowmelt may affect organisms in areas of late snowmelt most severely (Hein et al. 2014, Bowden et al. 2018, Wehner et al. 2023). Moreover, specialized species that present a narrow phenological range or host specialization like butterflies or parasitoid wasps will possibly be more vulnerable to trophic mismatch (Høye and Forchhammer 2008). As a contrasting example, body size can remain stable over decades (one arctic lycosid population: Høye et al. 2009). Global change may have a bigger impact on tropical insects because they are living close to their optimal temperature and they are relatively sensitive to temperature change (Deutsch et al. 2008). We also know that plasticity increases in an unpredictable environment (Fischer et al. 2011) and this raises again the issue of anthropic disturbance in stable environments.

Because of the variety of morphologies in arthropods, studies use diverse measures of body size and often did not compare or repeat their results within the same taxa. The results we present thus offer a rough overview of body size variation in arthropods. Multivariate analyses of body size provide a better approximation of a population’s body size (Caldas et al. 1996, Bidau and Martí 2007, Knapp and Knappová 2013). Ratios (Azevedo et al. 1998), allometry (Bidau and Martí 2007) or Geometric Morphometrics (GM) (Hernández et al. 2011) can further take shape into account. These methods require however a large sampling and analyzing effort, difficult to implement in large-scaled analyses. For future meta-analyses to be richer and more precise, open-source databases (Gallagher et al. 2020) and standardized traits measurements in arthropods (Moretti et al. 2017) and spiders (Lowe et al. 2020, Pekár et al. 2021)—including habitat and life-history traits—are essential. We plead for integrating more life history traits in monitoring and predicting research, which is increasingly easy due to the development of worldwide databases and mechanistic species distribution models, respectively.

## Conclusions

1. In arthropods and in spiders specifically, we argue that life-history traits such as voltinism, development, mobility and dispersal ability, guild—e.g. wanderer vs. web-building—are explanatory variables that influence body size patterns along macroecological gradients.
2. Multivariate assessment of body size is often lacking, although essential to grasp its variation across macroecological gradients. As an example, separate measurement of female and male body size can reveal sexual size dimorphism (SSD), an important trait to consider in spiders.
3. Abiotic factors such as microclimate, habitat are also key factors influencing body size variation, although they are not often reported.
4. In addition, we emphasize the need for future studies to use standardized measurements and to publish them in public databases to allow for comparable and more complete analyses.
5. Many rules have been set to fit body size trends, without however explaining the underlying patterns that drive body size clines (Meiri 2011). This review gathers empirical data about body size variation along elevational and latitudinal gradients in arthropods, providing a step forward to better understanding of underlying mechanisms driving body size variation.
6. The intraspecific study of body size through space and time is key in ecology, as it is driven by physiological and adaptive mechanisms, and addresses conservation and invasion issues (Violle et al. 2007, Thomas 2010, Wills et al. 2014, Bertelsmeier 2017, Austin and Dunlap 2019). However, most studies solely report correlations of phenotypic variation with macroecological gradients. We found many studies investigating body size variation along macroecological gradients, but we still know little about the causes and consequences of variation in life history trait plasticity in the wild, or how natural selection operates on plasticity (Pigliucci 2005, Stillwell 2010). A few studies combined body size measurements with common-garden experiments or genetic data on arthropods collected on macroecological gradients (Bulgarella et al. 2015, Sandoval-Arango et al. 2020, Yadav et al. 2021), but more of these studies are needed to address the mechanisms underlying body size variation along macroecological gradients.

## Acknowledgments

JP was supported by “BOOST ERC” OPALE and by the SAD “PEPPS” (both by the Région Bretagne).

## Supplementary Data

Datasets and R scripts used for both body size and SSD reviewing analysis are available in the Figshare repository at https://doi.org/10.6084/m9.figshare.27118986.v2.

## Tables

**Table 1:** The eight essential dichotomies implied by the study of body size variation in spiders in ecological and evolutionary research. The choices appear in bold.

| Restrictions for this study |  |
| --- | --- |
| Due to prior choices |  |
| 1. | **Ectotherms** vs. endotherms |
| 2. | **Intraspecific** vs interspecific variation |
| 3. | **Structural body size** vs. condition-dependent body size (see Box1) |
| Due to lack of studies |  |
| 4. | **Field** vs. experiments |
| 5. | **Bergmann's rule** vs. Temperature-Size Rule |
| 6. | **Macro** vs. microecological/spatial gradients |
| Possible comparisons within our field study |  |
| 7. | Direct vs. indirect development |
| 8. | Wandering vs. web-building spiders |

**Table 2:**
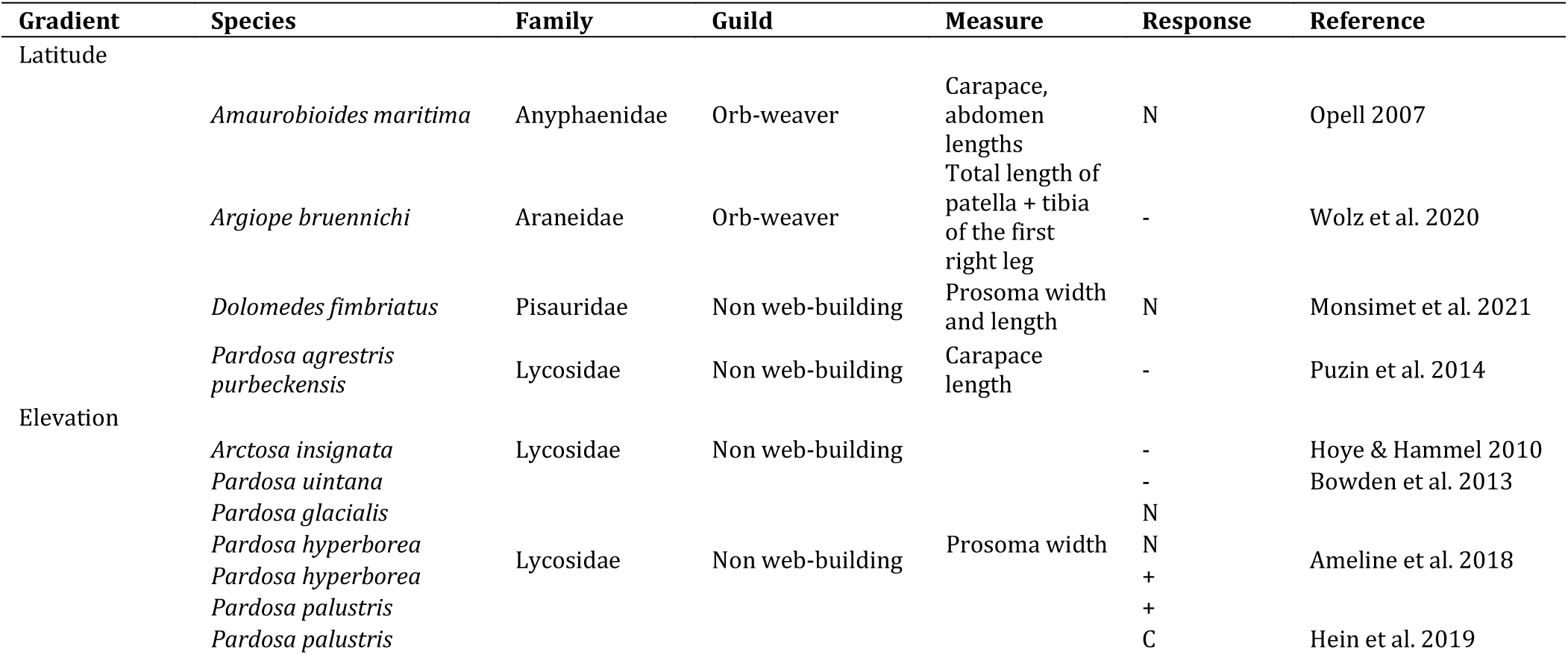
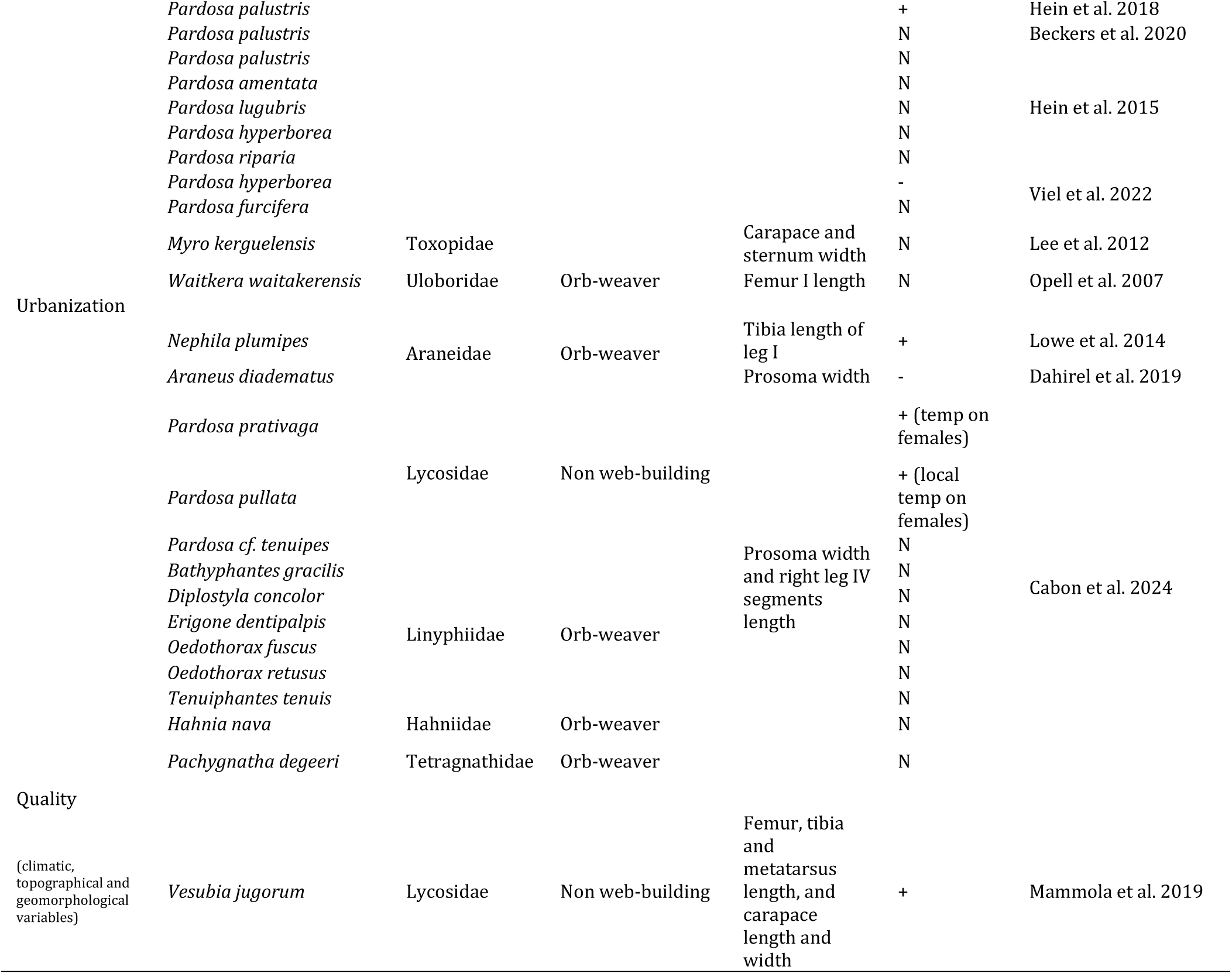
Body size variations in response to macroecological gradients in spiders. Response can be negative (-), positive (+), curvilinear (C), humped-shaped (H), none (N), sawtooth (S) or U-shaped (U). The full table of body size variation found in all arthropods is given in Supplementary Table S1 and at https://doi.org/10.6084/m9.figshare.27118986.v2. References: (Opell et al. 2007, Høye and Hammel 2010, Lee et al. 2012, Bowden et al. 2013, Lowe et al. 2014, Puzin et al. 2014, Hein et al. 2015, 2018, 2019, Ameline et al. 2018, Dahirel et al. 2019, Mammola et al. 2019, Beckers et al. 2020, Wolz et al. 2020, Monsimet et al. 2021, Viel et al. 2022, Cabon et al. 2024).

**Table 3:**
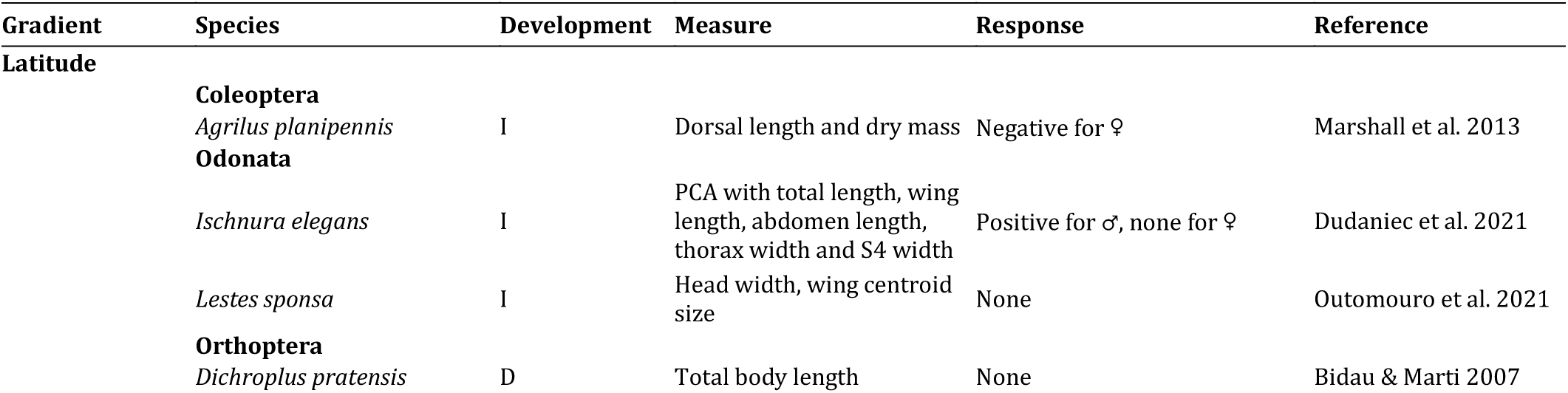
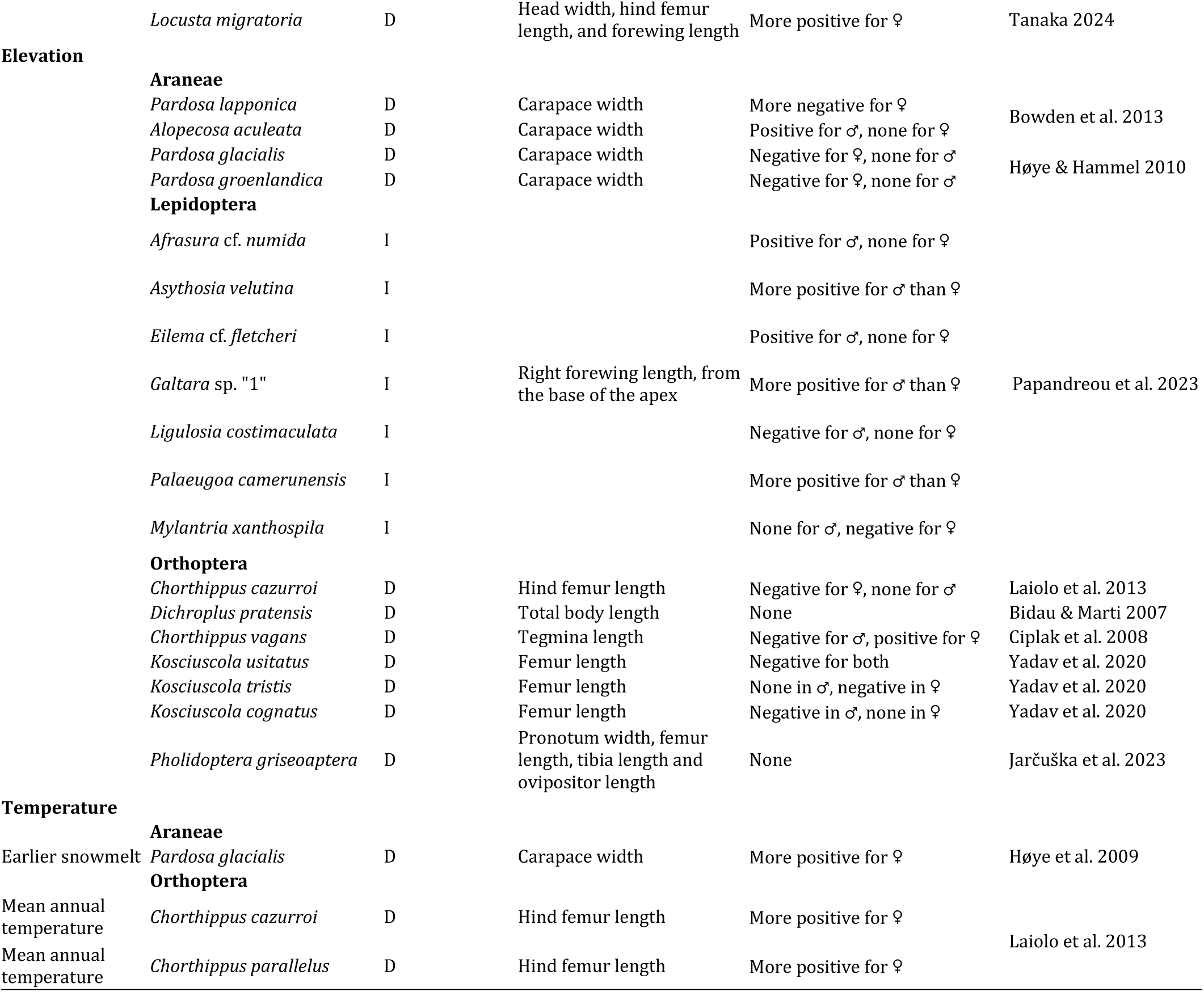
Sexual size dimorphism (SSD) variation along macroecological gradients in arthropods: D: direct, I: indirect development. References: (Bidau and Martí 2007, Ciplak et al. 2008, Høye et al. 2009, Høye and Hammel 2010, Bowden et al. 2013, Laiolo et al. 2013, Marshall et al. 2013, Yadav et al. 2020, Outomuro et al. 2021, Dudaniec et al. 2022, Jarčuška et al. 2023, Papandreou et al. 2023, Tanaka 2024).

