## Supplementary for "Spiders bring new insight into the eco-evolutionary drivers of body size variation and sexual size dimorphism in arthropods"

**Table S1:** Body size variations in response to latitudinal and elevational gradients. Response can be negative (-), positive (+), curvilinear (C), humped-shaped (H), none (N), sawtooth (S) or U-shaped (U). References with the "§", "£" and "*" symbols were listed in (Chown and Gaston 2010), (Blanckenhorn and Demont 2004) and (de Jong and Bochdanovits 2003), respectively. The excel version of this table is available at DOI: 10.6084/m9.figshare.27118986.

| Gradient | Group | Species | Development | Body size measure | Response | Reference |
|---|---|---|---|---|---|---|
| Elevation | | | | | | |
| | Araneae | *Arctosa insignata* | D | Carapace width | - | Hoye & Hammel 2010 |
| | | *Myro kerguelensis* | D | Carapace and sternum width | N | Lee et al. 2011 |
| | | *Pardosa amentata* | D | Pronotum width | N | Hein et al. 2015 |
| | | *Pardosa glacialis* | D | Prosoma width | N | |
| | | *Pardosa hyperborea* | D | Prosoma width | N | Ameline et al. 2018 |
| | | *Pardosa hyperborea* | D | Prosoma width | + | |
| | | *Pardosa hyperborea* | D | Pronotum width | N | Hein et al. 2015 |
| | | *Pardosa hyperborea* | D | Pronotum width | - | Viel et al. 2022 |
| | | *Pardosa furcifera* | D | Pronotum width | N | Viel et al. 2022 |
| | | *Pardosa lugubris* | D | Pronotum width | N | Hein et al. 2015 |
| | | *Pardosa palustris* | D | Prosoma width | + | Ameline et al. 2018 |
| | | *Pardosa palustris* | D | Pronotum width | N | Beckers et al. 2020 |
| | | *Pardosa palustris* | D | Pronotum width | N | Hein et al. 2015 |
| | | *Pardosa palustris* | D | Prosoma width | + | Hein et al. 2018 |
| | | *Pardosa palustris* | D | Prosoma width | C | Hein et al. 2019 |
| | | *Pardosa riparia* | D | Pronotum width | N | Hein et al. 2015 |
| | | *Pardosa uintana* | D | Carapace width | - | Bowden et al. 2013 |
| | | *Waitkera waitakerensis* | D | Femur I length | N | Opell et al. 2007 |
| | Cladocera | *Daphnia magna* | D | Head length, body width and length | + | Ma et al. 2020 |
| | Coleoptera | *Acanthoscelides macrophtalmus* | I | Elytron length and width, pronotum width | N | Haga and Rossi 2016 |
| | | *Adesmia metallica* | I | Elytron length | - | Krasnov et al. 1996§ |
| | | *Agrilus planipennis* | I | Body length | N | Nalepa et al. 2023 |
| | | *Amara alpina* | I | Pronotum width | - | Beckers et al. 2020 |
| | | *Amblystogenium pacificum* | I | Body size index from: pronotum length and width, femur and third left leg length, elytra width and length | H | Espel et al. 2023 |
| | | *Bothrometopus brevis* | I | Body length | N | |
| | | *Bothrometopus elongtaus* | I | Body length | + | |
| | | *Bothrometopus gracilipes* | I | Body length | - | Chown & Klok 2003§ |
| | | *Bothrometopus parvulus* | I | Body length | + | |
| | | *Canonopsis sericeus* | I | Body length | - | |
| | | *Canthon rutilans cyanescens* | I | Body length (clypeus to pygidium) | - | Oliveira de Alcântara et al. 2023 |
| | | *Carabus auronitens auronitens* | I | Elytron length | - | Baranovská et al. 2019 |
| | | *Carabus auronitens escheri* | I | Elytron length | N | |
| | | *Carabus exaratus* | I | Centroid size from geometric morphometrics | U | Benítez et al. 2023 |
| | | *Carabus japonicus* | I | Body length | - | Okuzaki and Sota 2017 |
| | | *Carabus linnei* | I | Elytron length | - | |
| | | *Carabus sylvestris sylvestris* | I | Elytron length | N | Baranovská et al. 2019 |
| | | *Carabus sylvestris transsylvanicus* | I | Elytron length | N | |
| | | *Carabus tosanus* | I | Body length | - | Ikeda et al. 2012 |

| | | | | | |
|---|---|---|---|---|---|
| | *Cephaloleia belti* | I | Body length, from the clypeus to the tip of the pygidium | + | González-Tokman et al 2019 |
| | *Chelobasis bicolor* | I | Body length, from the clypeus to the tip of the pygidium | + | |
| | *Dichotomius sericeus* | I | Body length (clypeus to pygidium) | N | Oliveira de Alcântara et al. 2023 |
| | *Ectemnorhinus marioni* | I | Body length | + | |
| | *Ectemnorhinus similis* | I | Body length | + | Chown & Klok 2003§ |
| | *Ectemnorhinus viridis* | I | Body length | - | |
| | *Erodius edomitus* | I | Elytron length | - | Krasnov et al. 1996§ |
| | *Lissorhoptrus oryzophilus* | I | Length of pronotum + elytra | + | Huang et al. 2018 |
| | *Nicrophorus investigator* | I | Elytron length | + | Smith et al. 2000§ |
| | *Onthophagus proteus* | I | Horn length, pronotum width | + | Stanbrook et al. 2021 |
| | *Onthophagus proteus* | I | Pronotal length, elytron length and width, abdomen depth and body length | N | Stanbrook et al. 2021 |
| | *Pimelia laevigata costipennis* | I | Elytra width and length | + | López et al 2021 |
| | *Pterostichus burmeisteri* | I | Elytron length | N | Baranovská et al. 2019 |
| | *Pterostichus melanarius* | I | Elytron length | - | Baranovská et al. 2019 |
| | *Pterostichus melanarius* | I | Elytron length | S | Luzyanin et al. 2022 |
| | *Pterostichus montanus* | I | Length and width of elytra, pronotum and head | N | Sukhodolskaya and Ananina 2017 |
| | *Pterostichus montanus* | I | Length and width of elytra, pronotum and head | + | Sukhodolskaya et al. 2021 |
| | *Pterostichus pilosus* | I | Elytron length | - | Baranovská et al. 2019 |
| | *Sepidium dathan* | I | Elytron length | + | Krasnov et al. 1996§ |
| | *Silpha carinata* | I | Elytron and pronotum width | - | Baranovská and Knapp 2018 |
| | *Zophosis complanata* | I | Elytron length | - | Krasnov et al. 1996§ |
| | *Zygogramma bicolorata* | I | Body length, from tip of head to end of elytra | + | Bhusal et al. 2019 |
| Copepoda | *Arctodiaptomus salinus* | I | Total length | - | Anufriieva and Shadrin 2014 |
| Diptera | *Arcynopteryx dicroa* | I | Body length | - | Loskutova and Zhiltzova 2017 |
| | *Drosophila immigrans* | I | Head width and body length | H | Fartyal et al. 2017 |
| | *Drosophila melanogaster* | I | Thorax length | + | Klepsatel et al. 2014 |
| | *Drosophila melanogaster* | I | fresh weight, wing length, wing area or thorax length | + | Fabian et al. 2015 |
| | *Drosophila nepalensis* | I | Head width and body length | H | Fartyal et al. 2017 |
| | *Drosophila repleta* | I | Head width and body length | H | Fartyal et al. 2017 |
| | *Scaptomyza himalayana* | I | Head width and body length | H | Fartyal et al. 2017 |
| | *Zaprionus grandis* | I | Head width and body length | H | Fartyal et al. 2017 |
| Ephemeroptera | *Drunella doddsii* | I | Body length (anterior end of the head to posterior of the abdomen) | N | McCarty et al. 2022 |
| | *Andesiops torrens* | I | Body length: | - | |
| | *Massartelopsis irarrazavali* | I | clypeus to tip of | N | Rendoll-Cárcamo et al. 2023 |
| | *Meridialaris chiloeensis* | I | last abdominal | + | |

| | | | | | |
|---|---|---|---|---|---|
| | *Metamonius anceps* | I | segment (excluding cerci) | + | |
| | *Ephemerella infrequens* | I | Body length (anterior end of the head to posterior of the abdomen) | - | McCarty et al. 2022 |
| Hemiptera | *Diaphorina citri* | D | Wing, thorax and antenna measurments | + | Pérez-Valencia and Moya-Raygoza 2015 |
| | *Laodelphax striatella* | D | Body, forewing, vertex, head, pronotum, mesonotum width and length | + | Karavin & Zeybekoglu 2023 |
| Hymenoptera | *Eoanthidium insulare* | I | Forewing area | N | Kasparek et al. 2024 |
| | *Bombus vancouverensis* | I | Forewing area | N | Lozier et al. 2021 |
| | *Bombus vosnesenskii* | I | Forewing area | + | Lozier et al. 2021 |
| | *Atta cephalotes* | I | Various head, mandible, thorax and leg measurments | H | Sandoval-Arango et al. 2020 |
| | *Bombus humilis* | I | Inter-tegular distance | + | Massa et al 2024 |
| | *Bombus wurflenii* | I | | + | |
| | *Bombus terrestris* | I | | - | |
| | *Bombus lapidarius* | I | | N | |
| | *Bombus pascuorum* | I | | N | |
| | *Cotesia flavipes* | I | mesosoma length | - | Lozano-Morales et al. 2024 |
| | *Cotesia flavipes* | I | | N | |
| | *Diglyphus isaea* | I | Thorax width, forewing length | + | Xi et al. 2024 |
| | *Eulaema nigrita* | I | Wing area | - | Vaca-Sánchez et al. 2023 |
| | *Ectomomyrmex javanus* | I | maximum head length, Weber's length: mesosoma length (body size), hind femur length | + | He et al. 2024 |
| | *Formica selysi* | I | Head width | + | Purcell et al. 2016 |
| | *Halictus* sp1a | | Intertegular distance | N | Osorio-Canadas et al. 2022 |
| | *Lasioglossum (Dialictis)* sp1 | I | | N | |
| | *Lasioglossum (Lasioglossum)* sp1 | I | | N | |
| | *Leptothorax acervorum* | I | maximum cephalic width (CW) and mesosoma length (ML) | + | Bernadou et al. 2016 |
| | *Macrotera* sp1 | I | Intertegular distance | N | Osorio-Canadas et al. 2022 |
| | *Odontoponera transversa* | I | maximum head length, Weber's length: mesosoma length (body size), hind femur length | + | He et al. 2024 |
| | *Pachyneuron aphidis* | I | Thorax width, forewing length | + | Xi et al. 2024 |
| | *Polistes annularis* | I | Multivariate PCA with head width, thorax length and forewing length | - | Miller and Sheehan 2021 |
| | *Polistes apachus* | I | | - | |
| | *Polistes aurifer* | I | | + | |
| | *Polistes bahamensis* | I | | + | |
| | *Polistes bellicosus* | I | | N | |
| | *Polistes carolina* | I | | - | |
| | *Polistes comanchus* | I | | N | |
| | *Polistes dominula* | I | | N | |
| | *Polistes dorsalis* | I | | + | |
| | *Polistes exclamans* | I | | + | |
| | *Polistes flavus* | I | | + | |
| | *Polistes fuscatus* | I | | - | |
| | *Polistes metricus* | I | | - | |

| | | | | | |
|---|---|---|---|---|---|
| | *Tetragonula biroi* | I | Body length and forewing length | + | Prastiyo et al. 2024 |
| Lepidoptera | *Danaus chrysippus* | I | | + | |
| | *Danaus genutia* | I | | + | |
| | *Vanessa indica* | I | Wing span, forewing length and width, wing surface area | + | Verma & Arya 2023 |
| | *Catopsilia pomona* | I | | - | |
| | *Pieris brassicae* | I | | - | |
| | *Papilio polytes* | I | | - | |
| | *Apodemia mormo* | I | | N | |
| | *Feniseca tarquinius* | I | | N | |
| | *Agriades glandon* | I | | N | |
| | *Glaucopsyche lygdamus* | I | | N | |
| | *Glaucopsyche piasus* | I | Forewing size | N | Merwin et al. 2022 |
| | *Satyrium edwardsii* | I | | N | |
| | *Callophrys irus* | I | | N | |
| | *Callophrys niphon* | I | | N | |
| | *Callophrys gryneus* | I | | N | |
| | *Afrasura* cf. *numida* | I | | + | |
| | *Afrasura* cf. *peripherica* | I | | + | |
| | *Anapisa* sp. "2" | I | | + | |
| | *Asura* cf. *craigii* | I | | + | |
| | *Asythosia velutina* | I | | + | |
| | *Cyana* sp. "2" | I | | - | |
| | *Eilema* cf. *fletcheri* | I | Right forewing length, from the base of the apex | + | Papandreou et al. 2023 |
| | *Galtara* sp. "1" | I | | + | |
| | *Lamprosiella* sp. "1" | I | | N | |
| | *Ligulosia costimaculata* | I | | - | |
| | *Palaeugoa camerunensis* | I | | + | |
| | *Phasmatilema* sp. "1" | I | | N | |
| | *Rhipidarctia postrosea* | I | | + | |
| | *Erebia medusa* | I | Forewing length and width | N | Mikitová et al. 2022 |
| | *Euproctis* sp. "1" | I | | N | |
| | *Leucoma* sp. "7" | I | | N | |
| | *Lymantriinae* sp. "5" | I | | N | |
| | *Lymantriinae* sp. "54" | I | | + | |
| | *Lymantriinae* sp. "55" | I | | + | |
| | *Lymantriinae* sp. "58" | I | | + | |
| | *Lymantriinae* sp. "65" | I | | N | |
| | *Lymantriinae* sp. "75" | I | Right forewing length, from the base of the apex | + | Papandreou et al. 2023 |
| | *Mylantria xanthospila* | I | | N | |
| | *Stracena* cf. *bananae* | I | | N | |
| | *Desmeocraera* sp. "1" | I | | N | |
| | *Eurystaura* sp. "1" | I | | N | |
| | *Mallocampa audea* | I | | - | |
| | *Mariaeia spargatana* | I | | - | |
| | *Polyptychus nigriplaga* | I | | + | |
| Orthoptera | *Aeropedellus clavatus* | D | Femur length | - | Levy and Nufio 2015 |
| | *Camnula pellucida* | D | Femur length | + | Levy and Nufio 2015 |
| | *Chorthippus vagans* | D | body, tegmina, pronotum, hind femur | N | Ciplak et al. 2008 |
| | *Dichroplus pratensis* | D | total body length, femur 3, tibia 3, tegmina and pronotum lengths, and pronotum height | - | Bidau & Marti 2007a |
| | *Dichroplus vittatus* | D | mean and maximum body length | - | Bidau & Marti 2007b |
| | *Hemideina crassidens* | D | Tibia length | + | Bulgarella et al. 2015 |
| | *Hemideina maori* | D | Head length | + | Koning & Jamieson 2001[§] |
| | *Kosciuscola usitatus* | D | Femur length | - | Yadav et al. 2020 |
| | *Kosciuscola tristis* | D | Femur length | - | Yadav et al. 2020 |
| | *Kosciuscola cognatus* | D | Femur length | N | Yadav et al. 2020 |
| | *Locusta migratoria* | D | Head width, hind femur length, and forewing length | N | Tanaka 2024 |
| | *Melanoplus boulderensis* | D | Femur length | - | Levy and Nufio 2015 |
| | *Melanoplus sanguinipes* | D | Femur length | + | Levy and Nufio 2015 |

| | | | | | |
|---|---|---|---|---|---|
| | | *Melanoplus sanguinipes* | D | Mass | + | Rourke 2000§ |
| | | *Oedipoda miniata* | D | body, tegmina, pronotum, hind femur | - | Ciplak et al. 2008 |
| | | *Omocestus viridulus* | D | Hind femur length | - | Berner & Blanckenhorn 2006§ |
| | | *Pholidoptera griseoaptera* | D | Pronotum width, femur length, tibia length and ovipositor length | - | Jarčuška et al. 2023 |
| | | *Poecilimon birandi* | D | body, tegmina, pronotum, hind femur | - | Ciplak et al. 2008 |
| | | *Poecilimon veluchianus minor* | D | Pronotum and hind femur length | N | Eweleit and Reinhold 2014 |
| | | *Poecilimon veluchianus veluchianus* | D | Pronotum and hind femur length | - | Eweleit and Reinhold 2014 |
| | | *Pseudochorthippus parallelus* | D | Pronotum length | N | Köhler et al. 2017 |
| | | *Pseudochorthippus parallelus* | D | Postfemur length | - | Köhler et al. 2017 |
| | | *Sphenarium histrio* | D | Femur III length | - | Ramírez-Delgado & Cueva del Castillo 2022 |
| | | *Teleogryllus emma* | D | Head width | - | Masaki 1967§ |
| | | *Xanthippus corallipes* | D | Mass | - | Ashby 1997§ |
| | Ostracoda | *Ilyocypris bradyi* | D | Valve length and heigth | + | |
| | | *Heterocypris salina* | D | Valve heigth | + | |
| | | *Heterocypris salina* | D | Valve length | N | |
| | | *Heterocypris incongruens* | D | Valve length and heigth | + | |
| | | *Potamocypris fallax* | D | Valve length and heigth | + | Dalgakiran et al. 2020 |
| | | *Psychrodromus olivaceus* | D | Valve length and heigth | + | |
| | | *Ilyocypris inermis* | D | Valve length and heigth | N | |
| | | *Heterocypris reptans* | D | Valve length and heigth | N | |
| | Plectoptera | *Limnoperla jaffueli* | I | Body length: clypeus to tip of last abdominal segment (excluding cerci) | N | Rendoll-Cárcamo et al. 2023 |
| | | *Rhithroperla rossi* | I | | N | |
| | | *Senzilloides panguipulli* | I | | + | |
| | | *Stenoperla prasina* | I | Forewing length | N | Winterbourn et al. 2017 |
| | | *Stenoperla prasina* | I | Forewing length | + | Winterbourn et al. 2017 |
| | | *Udamocercia antarctica* | I | Body length: clypeus to tip of last abdominal segment (excluding cerci) | N | Rendoll-Cárcamo et al. 2023 |
| | Trichoptera | *Hydropsyche ambigua* | I | Male right anterior wing | + | Cogo et al 2020 |
| | | *Hydropsyche cockerelli* | I | Boday length (anterior end of the head to the beginning of the anal proleg) | S | McCarty et al. 2022 |
| | | *Hydropsyche siltalai* | I | Male right anterior wing | + | Cogo et al 2020 |
| | | *Rhyacophila adjuncta* | I | Male right anterior wing | + | Cogo et al 2020 |
| Latitude | Acari | *Laelaps clethrionomydis* | I | Length of dorsal and sternal shields | + | Korallo-Vinarskaya et al. 2015 |
| | Amphipoda | *Orchestoidea tuberculata* | D | Body length: rostrum tip to telson base | + | Jaramillo et al. 2017 |
| | Araneae | *Amaurobioides maritima* | D | Carapace, abdomen lengths | N | Opell 2010 |

| | | | | | |
|---|---|---|---|---|---|
| | *Argiope bruennichi* | D | total length of patella + tibia of the first right leg | - | Wolz et al. 2020 |
| | *Dolomedes fimbriatus* | D | Prosoma width and length | N | Monsimet et al. 2021 |
| | *Pardosa agrestris purbeckensis* | D | Carapace length | - | Puzin et al. 2014 |
| Blattodea | *Eupolyphaga sinensis* | D | body length, body width, pronotum width | S | Hu et al. 2012 |
| | *Reticulitermes speratus* | D | Head width | + | Morooka et al. 2024 |
| Chilopoda | *Strigamia maritima* | D | Body length | - | Hayden et al. 2012 |
| Coleoptera | *Acanthoscelides macrophtalmus* | I | Elytron length and width, pronotum width | N | Haga and Rossi 2016 |
| | *Acanthoscelides pallidipennis* | I | Elytral length | + | Sadakiyo & Ishihara 2012 |
| | *Agrilus planipennis* | I | Body length | S | Nalepa et al. 2023 |
| | *Carabus cancellatus* | I | | - | |
| | *Carabus granulatus* | I | Elytra length | - | Sukhodolskaya and Saveliev 2016 |
| | *Carabus hortensis* | I | | - | |
| | *Carabus nemoralis* | I | Elytron length | - | Blanckenhorn & Demont 2004[§] |
| | *Carabus nemoralis* | I | Elytron length | - | Krumbiegel 1936 (1932 ?)[£] |
| | *Ceroglossus chilensis* | I | Wing centroid size | - | Benítez et al. 2024 |
| | *Colaphellus bowringi* | I | Body weight | - | Tang et al. 2017 |
| | *Dendroctonus ponderosae* | I | Pronotal width | - | Bentz et al. 2001 |
| | *Dicaelus purpuratus* | I | length | - | Park 1949[§] |
| | *Lissorhoptrus oryzophilus* | I | Length of pronotum + elytra | + | Huang et al. 2018 |
| | *Paropsis atomaria* | I | protonum width | - | Schutze & Clarke 2008 |
| | *Phaleria testacea* | I | Head width, pronotal width, elytral length | N | Caldas et al. 1996 |
| | *Phyllotreta striolata* | I | Elytron length | + | Blanckenhorn & Demont 2004[§] |
| | *Phyllotreta striolata* | I | Elytron length | + | Masaki 1967[£] |
| | *Poecilus cupreus* | I | Elytra and head size | + | Sukhodol et al. 2017 |
| | *Poecilus cupreus* | I | Pronotum size | - | Sukhodol et al. 2017 |
| | *Poecilus cupreus* | I | Elytra length | N | Sukhodolskaya and Saveliev 2016 |
| | *Pterostichus melanarius* | I | Elytra length | S | Sukhodolskaya and Saveliev 2016 |
| | *Pterostichus melanarius* | I | Elytra length | S | Luzyanin et al. 2022 |
| | *Pterostichus niger* | I | Elytra length | - | Sukhodolskaya and Saveliev 2016 |
| | *Stator limbatus* | I | Elytron length/ width | + | Stillwell et al. 2007[§] |
| | *Tribolium castaneum* | I | | + | Matsumara et al. 2023 |
| Decapoda | *Emerita analoga* | I | Carapace length: rostrum tip to distal scoop | + | Jaramillo et al. 2017 |
| Diptera | *Aedes aegypti (L.)* | I | Wing length | U | Neoh et al. 2024 |
| | *Aedes sierrensis* | I | Wing length | + | Lyberger et al. 2024 |
| | *Arcynopteryx dicroa* | I | Body length | - | Loskutova and Zhiltzova 2017 |
| | *Bactrocera tryoni* | I | Intertegular length | + | Zhou et al. 2020 |
| | *Drosophila melanogaster* | I | Thorax length | + | Klepsatel et al. 2014 |
| | *Drosophila melanogaster* | I | | + | David 1975* |
| | *Drosophila melanogaster* | I | | + | Delpuech et al. 1995* |
| | *Drosophila melanogaster* | I | | + | Noach et al. 1996* |
| | *Drosophila melanogaster* | I | | + | Bochdanovits and de Jong 2003* |
| | *Drosophila melanogaster* | I | | + | Imasheva et al. 1994* |
| | *Drosophila melanogaster* | I | fresh weight, wing length, wing area or thorax length | + | Coyne and Beecham 1987* |
| | *Drosophila melanogaster* | I | | + | Bochdanovits and de Jong 2003* |
| | *Drosophila melanogaster* | I | | + | James et al. 1995, 1997* |
| | *Drosophila melanogaster* | I | | + | van't Land et al. 1999* |
| | *Drosophila melanogaster* | I | | + | Capy et al. 1993* |
| | *Drosophila melanogaster* | I | | + | Gilchrist and Partridge 1999* |
| | *Drosophila melanogaster* | I | | N | David et al. 1976* |

| | | | | | |
|---|---|---|---|---|---|
| | *Drosophila melanogaster* | I | | + | Fabian et al. 2015 |
| | *Drosophila melanogaster* | I | | + | |
| | *Scathophaga stercoraria* | I | Hind tibia length | H | Bauerfeind et al. 2018 |
| | *Scathophaga stercoraria* | I | Hind tibia length | N | Blanckenhorn et al 2018 |
| Hemiptera | *Feniseca tarquinius* | D | Wing centroid size | N | Carbajal-de-la-Fuente et al. 2024 |
| Hymenoptera | *Aphidius platensis* | I | Hind tibia length (right) | N | Alfaro-Tapia et al. 2021 |
| | *Eoanthidium insulare* | I | Forewing area | + | Kasparek et al. 2024 |
| | *Bombus vancouverensis* | I | Forewing area | + | Lozier et al. 2021 |
| | *Bombus vosnesenskii* | I | Forewing area | N | Lozier et al. 2021 |
| | *Leptothorax acervorum* | I | Thorax length | + | Heinze et al. 1998[§], 2003[§]; A. Ruschinger, pers. comm.[£] |
| | *Myrmica rubra* | I | Mass | C | Elmes et al. 1999[§] |
| | *Polistes annularis* | I | Multivariate PCA with head width, thorax length and forewing length | - | Miller and Sheehan 2021 |
| | *Polistes apachus* | I | | - | |
| | *Polistes aurifer* | I | | N | |
| | *Polistes bahamensis* | I | | N | |
| | *Polistes bellicosus* | I | | + | |
| | *Polistes carolina* | I | | - | |
| | *Polistes comanchus* | I | | + | |
| | *Polistes dominula* | I | | - | |
| | *Polistes dorsalis* | I | | + | |
| | *Polistes exclamans* | I | | + | |
| | *Polistes flavus* | I | | - | |
| | *Polistes fuscatus* | I | | - | |
| | *Polistes metricus* | I | | - | |
| Isopoda | *Excirolana braziliensis* | D | Body length: rostrum tip to telson tip | + | Jaramillo et al. 2017 |
| | *Excirolana hirsuticauda* | D | Body length: rostrum tip to telson tip | + | Jaramillo et al. 2017 |
| Lepidoptera | *Apodemia mormo* | I | Forewing size | N | Merwin et al. 2022 |
| | *Feniseca tarquinius* | I | | N | |
| | *Agriades glandon* | I | | - | |
| | *Glaucopsyche lygdamus* | I | | N | |
| | *Glaucopsyche piasus* | I | | N | |
| | *Satyrium edwardsii* | I | | - | |
| | *Callophrys irus* | I | | - | |
| | *Callophrys niphon* | I | | - | |
| | *Callophrys gryneus* | I | | - | |
| Odonata | *Calopteryx maculata* | I | Only males, PC1 of fore wing and hind wing lengths and the length of the hind tibia | U | Hassal 2013 |
| | *Calopteryx splendens* | I | Head width and thorax length | + | Golab et al. 2022 |
| | *Erythromma viridulum* | I | PC1 of 7 size traits from head, wing, tibia, thorax and abdomen | + | Hassall et al. 2014 |
| | *Ischnura elegans* | I | PCA with total length, wing length, abdomen length, thorax width and S4 width | + | Dudaniec et al. 2021 |
| | *Lestes sponsa* | I | Head width | - | Sniegula et al. 2016 |
| | *Lestes sponsa* | I | Head width, wing centroid size | U | Outomouro et al. 2021 |
| | *Lestes sponsa* | I | Wing size | - | Yildirim et al. 2024 |
| Opiliones | *Leiobunum japonicum* | D | First leg femur | - | Tsurusaki & Okuda 2023 |
| Orthoptera | *Acheta pennsylvanicus* | D | Body length | - | Bigelow 1962[£] |
| | *Acheta veletis* | D | Body length | - | Alexander & Bigelow 1960[§] |

| | | | | | |
|---|---|---|---|---|---|
| | *Acheta veletis* | D | Body length | - | Alexander & Bigelow 1960[£] |
| | *Allonemobius fasciatus* | D | Femur length | S | Mousseau & Roff 1989[§] |
| | *Allonemobius socius* | D | Femur length | - | Mousseau & Roff 1989[£]; Bradford & Roff 1993[£] |
| | *Caledia captiva* | D | Pronotum length | - | Groeters & Shaw 1996[§] |
| | *Chorthippus brunneus* | D | Mass | - | Telfer & Hassall 1999[§] |
| | *Chorthippus brunneus* | D | Mass | - | Telfer & Hassall 1999[£] |
| | *Conocephalus spartinae* | D | Hind tibia length | - | Wason & Pennings 2008 |
| | *Decticus albifrons* | D | Body length | N | Samways & Harz 1982 |
| | *Decticus verrucivorus* | D | Body length | - | Samways & Harz 1982 |
| | *Dichroplus pratensis* | D | Total body length, femur 3, tibia 3, tegmina and pronotum lengths, and pronotum height | - | Bidau & Marti 2007a |
| | *Dichroplus vittatus* | D | Mean and maximum body length | - | Bidau & Marti 2007b |
| | *Locusta migratoria* | D | Head width, hind femur length, and forewing length | S | Tanaka 2024 |
| | *Melanoplus femurrubrum* | D | Hind femur length | - | Parsons and Joern 2014 |
| | *Orchelimum fidicinium* | D | Tibia length, body length | - | Ho et al. 2010 |
| | *Orchelimum fidicinium* | D | Hind tibia length | - | Wason & Pennings 2008 |
| | *Pteronemobius fascipes* | D | Head width | - | Masaki 1972[£] |
| | *Pteronemobius taprobanensis* | D | Head width | S | Masaki 1978[§] |
| | *Teleogryllus emma* | D | Head width | - | Masaki 1967[§] |
| | *Teleogryllus yezoemma* | D | Head width | - | Ohmachi & Masaki 1964[£] |
| | *Velarifictorus micado* | D | Head width | - | Zeng and Zhu 2014 |
| Ostracoda | *Cyprideis torosa* | D | Valve size | + | Wrozyna et al. 2022 |
| Plectoptera | *Stenoperla prasina* | I | Forewing length | N | Winterbourn et al. 2017 |
| | *Stenoperla prasina* | I | Forewing length | + | Winterbourn et al. 2017 |

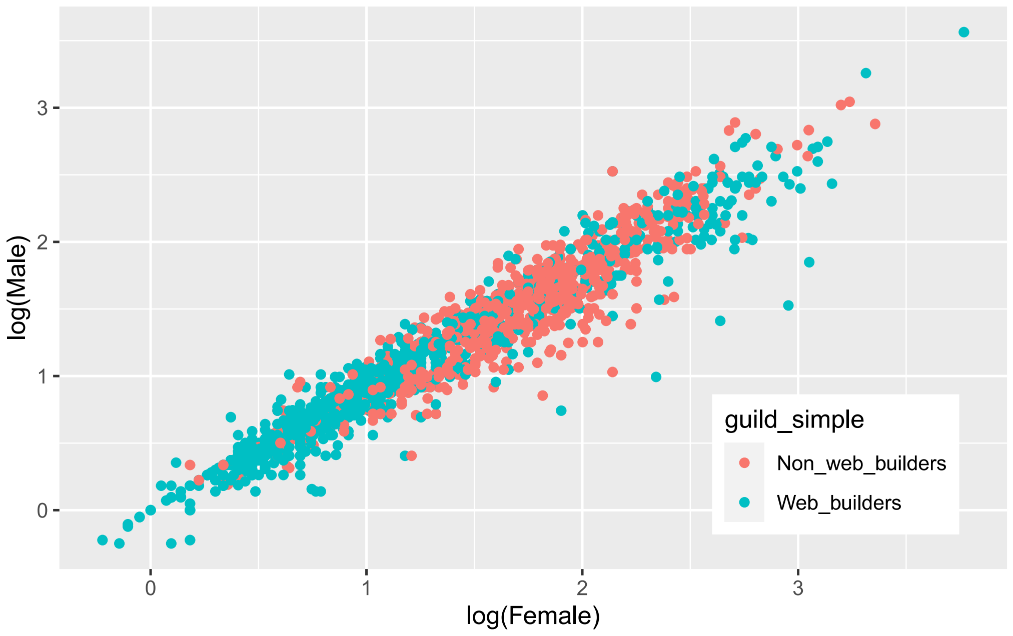


**Figure S1** Sexual size dimorphism in non web-building and in web-building spiders.

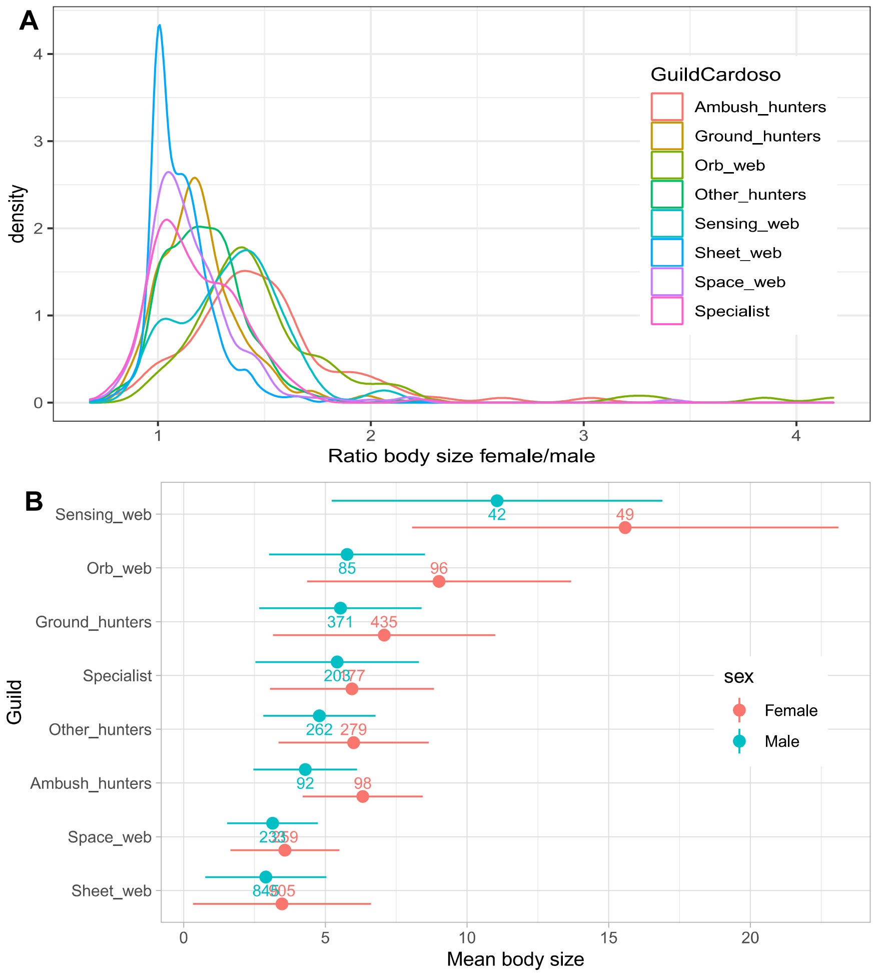


**Figure S2** Sexual size dimorphism in different spider guilds.

### References

**Table S1**

(Samways and Harz 1982; Caldas et al. 1996; Krasnov et al. 1996; Smith et al. 2000; Bentz et al. 2001; Chown and Klok 2003; Heinze et al. 2003; de Jong and Bochdanovits 2003; Blanckenhorn and Demont 2004; Bidau and Martí 2007; Opell et al. 2007; Stillwell et al. 2007; Bidau and Martí 2008a; Bidau and Martí 2008b; Ciplak et al. 2008; Schutze and Clarke 2008; Wason and Pennings 2008; Ho et al. 2010; Høye and Hammel 2010; Hayden et al. 2012; Hu et al. 2012; Ikeda et al. 2012; Lee et al. 2012; Sadakiyo and Ishihara 2012; Bowden et al. 2013; Hassall 2013; Anufriieva and Shadrin 2014; Eweleit and Reinhold 2014; Hassall et al. 2014; Klepsatel et al. 2014; Parsons and Joern 2014; Puzin et al. 2014; Zeng and Zhu 2014; Bulgarella et al. 2015; Fabian et al. 2015; Hein et al. 2015; Korallo-Vinarskaya et al. 2015; Levy and Nufio 2015; Pérez-Valencia and Moya-Raygoza 2015; Bernadou et al. 2016; Haga and Rossi 2016; Loskutova and Zhiltzova 2016; Purcell et al. 2016; Sniegula et al. 2016; Sukhodolskaya and Saveliev 2016; Jaramillo et al. 2017; Köhler et al. 2017; Okuzaki and Sota 2017; Sukhodol et al. 2017; Sukhodolskaya and Ananina 2017; Tang et al. 2017; Winterbourn et al. 2017; Ameline et al. 2018; Baranovská and Knapp 2018; Bauerfeind et al. 2018; Blanckenhorn et al. 2018; Fartyal et al. 2018; Hein et al. 2018; Huang et al. 2018; Baranovská et al. 2019; Bhusal et al. 2019; González-Tokman et al. 2019; Hein et al. 2019; Beckers et al. 2020; Cogo et al. 2020; Dalgakıran et al. 2020; Ma et al. 2020; Sandoval-Arango et al. 2020; Wolz et al. 2020; Yadav et al. 2020; Zhou et al. 2020; López et al. 2021; Lozier et al. 2021; Miller and Sheehan 2021; Miller and Sheehan 2021; Monsimet et al. 2021; Outomuro et al. 2021; Stanbrook et al. 2021; Sukhodolskaya et al. 2021; Alfaro-Tapia et al. 2022; Dudaniec et al. 2022; Golab et al. 2022; Luzyanin et al. 2022; McCarty et al. 2022; Merwin et al. 2022; Mikitová et al. 2022; Osorio-Canadas et al. 2022; Ramírez-Delgado and Cueva Del Castillo 2022; Viel et al. 2022; Wrozyna et al. 2022; Benítez et al. 2023; Espel et al. 2023; Jarčuška et al. 2023; Karavin and Zeybekoğlu 2023; Oliveira De Alcântara et al. 2023; Oliveira De Alcântara et al. 2023; Papandreou et al. 2023; Rendoll-Cárcamo et al. 2023; Tsurusaki and Okuda 2023; Vaca-Sánchez et al. 2023; Verma and Arya 2023; Benítez et al. 2024; Carbajal-de-la-Fuente et al. 2024; He et al. 2024; Kasparek et al. 2024; Lozano-Morales et al. 2024; Lyberger et al. 2024; Massa et al. 2024; Morooka et al. 2024; Nalepa et al. 2024; Neoh et al. 2024; Prastiyo et al. 2024; Tanaka 2024; Xi et al. 2024; Yildirim et al. 2024)
